# Chemical screen of Arabidopsis zygote and proteomics in tobacco BY-2 cells identify general plant cell division inhibitors

**DOI:** 10.1101/2022.04.28.489799

**Authors:** Yusuke Kimata, Moé Yamada, Takashi Murata, Keiko Kuwata, Ayato Sato, Takamasa Suzuki, Daisuke Kurihara, Mitsuyasu Hasebe, Tetsuya Higashiyama, Minako Ueda

**Author notes:** **Corresponding author:** Minako Ueda. **Author Contributions:** Y.K., A.S., D.K., T.H., and M.U. designed the research; Y.K., M.Y., T.M., K.K., T.S., D.K., M.H., and M.U. performed the research; Y.K., K.K., T.S., and M.U. analyzed the data; and Y.K., M.Y., T.M., D.K., and M.U. wrote the manuscript.

## Abstract

Cell division is essential for growth and development and involves events such as spindle assembly, chromosome separation, and cell plate formation. In plants, the tools used to control these events at the desired time are still poor because the genetic approach is ineffective owing to a high redundancy and lethality, as well as harmful side effects. Accordingly, we screened cell division-affecting compounds, with a focus on *Arabidopsis thaliana* zygotes, which individually develop in maternal ovules; the cell division was reliably traceable without time-lapse observations. We then identified the target events of the identified compounds using tobacco BY-2 cells for live-cell imaging and proteomics. As a result, we isolated two compounds, PD-180970 and PP2. PD-180970 disrupts microtubule (MT) organization and, thus, nuclear separation, presumably by inhibiting MT-associated proteins (MAP70). PP2 affected class II Kinesin-12 localization at the phragmoplast emerging site and impaired cytokinesis. Moreover, neither chemical caused irreversible damage to viability but they were effective in multiple plant species such as cucumber (*Cucumis sativus*) and moss (*Physcomitrium patens*). We propose that the combination of chemical screening based on Arabidopsis zygotes and target event specification focusing on tobacco BY-2 cells can be used to effectively identify novel tools and transiently control specific cell division events that are conserved in diverse plant species.

## Introduction

In multicellular organisms, growth and pattern formation depend on accurate cell division. Plant cell division requires intracellular events such as chromosome condensation and spindle assembly for nuclear division as well as the formation of a preprophase band (PPB) and phragmoplast for cytokinesis (1). Manipulating specific events at the desired time is crucial for controlling development and analyzing the cell division machinery. However, developing such tools is still challenging in plants because genetic manipulations do not work effectively owing to a high gene redundancy, mutant lethality, and various pleiotropic effects. For example, in *Arabidopsis thaliana* (Arabidopsis), multiple mutations in related cell division regulators cause gametophytic or embryonic lethality, whereas single mutants display no or minor defects (2, 3). In addition, mutants that fail to regulate microtubules (MT), a pivotal component of the spindle, PPB, and phragmoplasts, inhibit cell division as well as exhibit pleiotropic phenotypes such as affected stress responses and tissue specification (4).

To overcome redundancy, tools such as chemical inhibitors that bind to the conserved domains of homologous proteins are required to simultaneously impede the functions of related regulators (5). Furthermore, inhibitors can be applied at any time and therefore represent powerful tools that can ensure transient and stage-specific control as well as avoid lethality and side effects. Accordingly, in animal studies, the development of various specific inhibitors has greatly advanced our understanding of cell division mechanisms (6).

In plants, chemical effects on cell division control can be adequately monitored using time-lapse observations of the *Nicotiana tabacum* (tobacco) Bright Yellow-2 (BY-2) cell strain (7). BY-2 suspension cultures are suitable in performing intracellular and biochemical analyses such as high-speed live imaging of phragmoplast formation and purifications of phosphorylated proteins from synchronized cells (8, 9). However, in suspension cultures, cells at different division stages are mixed, and their positions in liquid media constantly change. Therefore, skilled professionals and specific devices for time-lapse observations are needed to trace individual cell states and thus determine the exact effects of the tested compounds on cell division.

Arabidopsis zygotes are single cells with thoroughly documented cell division time points (time courses) and patterns that can be easily identified in the individual ovules (10, 11). For example, it takes approximately 20 h after fertilization before the zygote commits to its first cell division, and the average time required for subsequent cell divisions is 7-9 h until the early globular embryo is formed (12). The distinct anatomy and regular cell division durations of the zygote make it an ideal platform to investigate cell division events and identify any morphological deviation. In addition, a reliable ovule cultivation system for time-lapse observations has been established to record the developmental time course of growing zygotes (12–14). Using high-resolution microscopy, the ovule cultivation system can be utilized to evaluate pharmacological effects on zygotic division and monitor intracellular dynamics (15–17).

In this study, we introduced an ovule cultivation system to screen compounds that affect zygotic cell division. The effects of the identified compounds on cell division were assessed using timelapse observations of tobacco BY-2 cells. Using mass spectrometry (MS)-based protein identification and assessments using various cell types and plant species, we identified two plant cell division inhibitors, PD-180970 and PP2, which generally block MT organization and phragmoplast formation, respectively, in diverse plant species.

## Results

### Identifying potent plant cell division inhibitors based on chemical screening of Arabidopsis zygote

In this study, we established an *in vitro* ovule cultivation system for chemical screening. To observe cell division, we used a dual-color marker that simultaneously labels the embryonic nuclei and plasma membranes (histone/PM) (12). Self-pollinated and zygote-containing ovules were isolated from the pistils and incubated in a culture media for two days (Fig. 1A). When cultivated with a control solvent (dimethyl sulfoxide, DMSO), all fluorescent-positive (living) embryos developed into the globular stage without causing morphological defects (Fig. 1B, DMSO), as indicated by the ratio of arrested and abnormal embryos (0%, n = 132). When cultivated with a known MT inhibitor, oryzalin, most zygotes did not divide (93%, n = 28; Fig. 1B, Oryzalin). Therefore, we concluded that this ovule cultivation system is capable of examining the inhibitory effects of the applied compounds and subsequently screened for potential plant cell division inhibitors.

**Figure 1.**
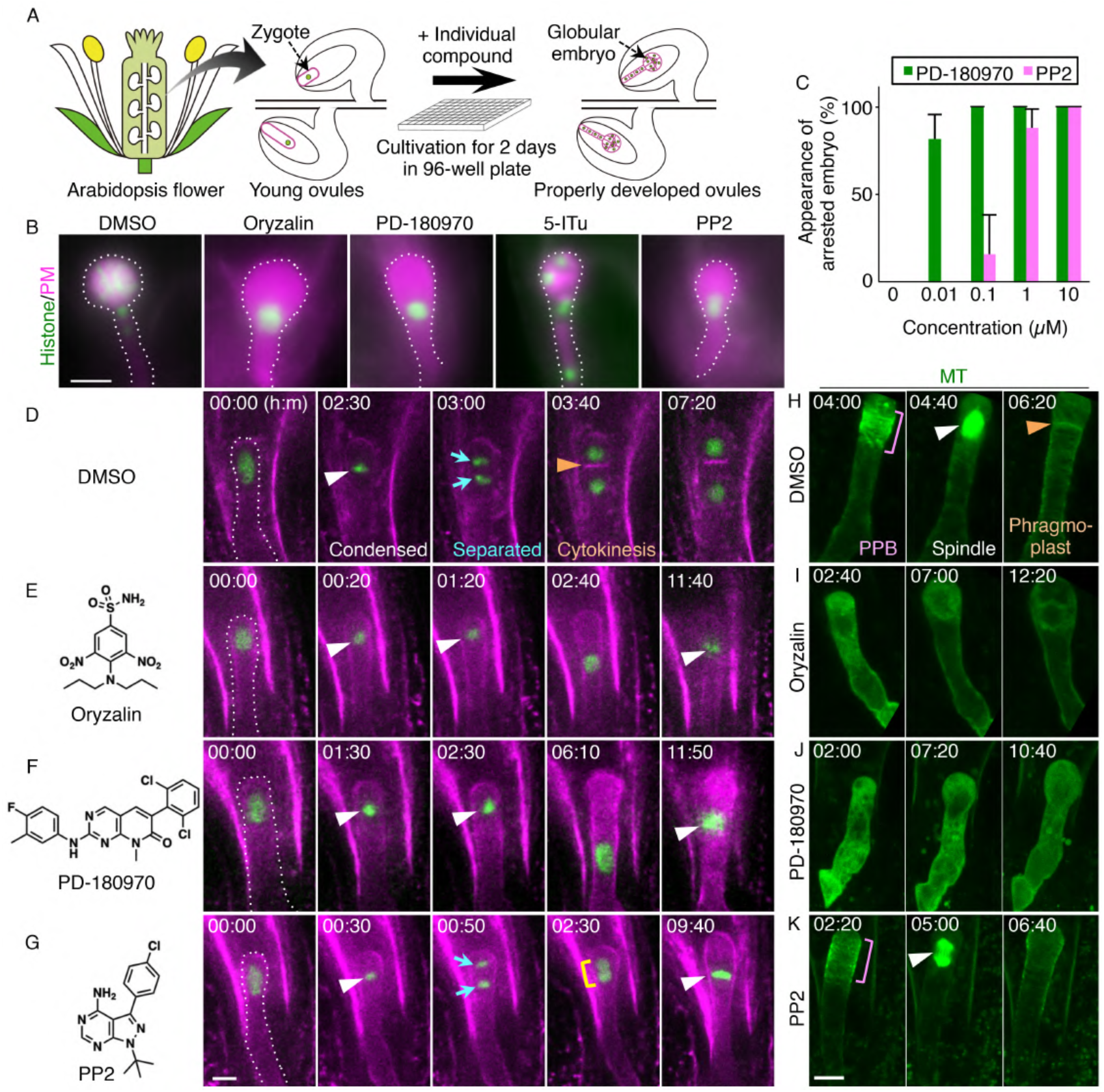
Chemical screening based on Arabidopsis zygote to identify cell division inhibitors. (A) Schematic procedure of chemical screening using *in vitro* ovule cultivation system. (B) Epi-fluorescent images of the embryos expressing histone/PM marker at 2 days after the application of indicated compounds. Embryos are outlined by dotted lines. (C) Dosage-dependent inhibition of Arabidopsis zygote over the concentrations of PD-180970 and PP2. Appearance of arrested embryos is the ratio of embryos with fewer cells compared to globular embryos among total living embryos at 2 days after the incubation. Error bars represent SD (n = 3). More than 52 ovules were counted for each test. (D-G) Time-lapse observation of the zygote expressing the histone/PM marker in the presence of the indicated compounds. The left drawings show the structures of respective compounds. Numbers indicate the time (h:min) from the first frame, and the starting zygotes are outlined by dotted lines. White and orange arrowheads indicate the spindle and newly formed cell plate, respectively. Cyan arrows and yellow rectangle show separated nuclei and two accompanied nuclei, respectively. (H-K) Time-lapse observation of MT alignment in Arabidopsis zygote in the presence of indicated compounds. Maximum intensity projection (MIP) images are shown, and numbers indicate the time (h:min) from the first frame. Magenta rectangles mark PPB formation in dividing zygotes. White and orange arrowheads indicate the spindle and phragmoplast, respectively. Scale bars: 30 μm (B) and 10 μm (D-K).

Two commercially available chemical libraries were selected for screening. The <u>L</u>ibrary <u>O</u>f <u>P</u>harmacologically <u>A</u>ctive <u>C</u>ompounds (LOPAC®) Pfizer library (Sigma-Aldrich) consists of 90 bioactive compounds whose targets have already been identified in animals. The SCREEN-WELL^®^ Kinase Inhibitor Library (Enzo Life Science) includes 80 reagents, each of which inhibits specific kinases in animals. Individual compounds were applied to cultivated ovules at a 10 μM concentration; this was because both libraries were pre-solubilized at 10 mM in DMSO, and DMSO exceeding 0.1% as the final concentration harms cell division in this cultivation system. Three antiproliferative compounds were identified with two technical replications: PD-180970 and 5-Iodotubercidin (5-ITu) from LOPAC® Pfizer, and PP2 from the SCREEN-WELL® Kinase Inhibitor (Fig. 1B). Zygotes cultivated with PD-180970 (100 %, n = 50) and PP2 (100 %, n = 78) showed severe cell division arrest as similar to those cultivated with oryzalin (Fig. 1B). Moreover, PD-180970 and PP2 were effective at concentrations of 10 nM and 1 μM, respectively (Fig. 1C). In contrast, 5-ITu only partially inhibited cell division and resulted in abnormal embryos with fewer cells compared to the DMSO-treated embryos (77%, n = 35, Fig. 1B). The 5-ITu compound has reportedly inhibited Haspin kinase in both animals and plants, and has cause chromosome misalignment during mitosis in tobacco BY-2 cells (18, 19). In contrast, PD-180970 and PP2 inhibit specific tyrosine kinases in animals, Bcr-Abl and Src, respectively (20, 21). However, their targets in plants were not predicted because these kinases have no homologs in plants. Therefore, we focused on PD-180970 and PP2 for further analyses.

### PD-180970 and PP2 respectively inhibit MT organization and phragmoplast formation

To obtain additional insights into the inhibitory mechanisms of PD-180970 and PP2, we examined the cellular dynamics of cultivated zygotes using a time-lapse observation system (Fig. 1D-G and Movie S1). In DMSO-treated zygotes, the zygotic nucleus was condensed and then separated. This was followed by cell plate formation, which is indicative of cytokinesis, resulting in two decondensed nuclei in separate cells (Fig. 1D). In oryzalin- and PD-180970-treated zygotes, the nucleus failed to separate and no cell plates were formed, resulting in one decondensed nucleus per cell (Fig. 1E and F). In PP2-treated zygotes, the zygotic nucleus condensed and separated normally, but no cell plate was formed, resulting in two accompanying nuclei or one fused nucleus in a cell (Fig. 1G).

Because MT pivotally influences both mitosis and cytokinesis, we performed time-lapse observations to examine the effects of PD-180970 and PP2 on MT markers in zygotes using two-photon excitation microscopy (2PEM) (Fig. 1H-K and Movie S2). The PPB, spindle, phragmoplast, and cortical MT arrays were clearly observed in DMSO-treated zygotes, as shown previously (Fig. 1H) (15). These MT structures were abolished in the oryzalin- and PD-180970-treated zygotes (Fig. 1I and J). In contrast, only phragmoplasts were lost in the PP2-treated zygotes (Fig. 1K). In addition, none of the compounds disrupted the longitudinal alignment of the other cytoskeleton, actin filaments (F-actin), following 1 h of treatment (Fig. S1A), whereas the same conditions caused a disruption of the F-actin array in the presence of an F-actin-specific inhibitor (15). These results suggest that PD-180970 and PP2 specifically affect MT alignment and phragmoplast formation, respectively.

To further assess whether the inhibitory effects were specific to Arabidopsis zygotes or general to other plants, we performed time-lapse observations on the BY–GTRC strain, in which MT and centromeric histones were labelled in tobacco BY-2 cells (MT/centromere; Fig. S1B and Movie S3) (22). The same effects observed in Arabidopsis zygotes were also observed in tobacco BY-2 cells, that is, PD-180970 and oryzalin inhibited nuclear division and PP2 blocked phragmoplast formation (Fig. S1B and Movie S3). Therefore, we concluded that these compounds targeted essential cell division regulators that are common in Arabidopsis zygotes and tobacco suspension-cultured cells.

### Protein identification utilizing BY-2 cells for PD-180970 and PP2

To investigate the inhibitory mechanisms of the compounds, we utilized the tobacco BY-2 cell system for subsequent proteomic analyses to identify the target proteins of PD-180970 and PP2. To obtain non-effective compounds as negative controls for proteomic analyses, we examined PD-180970 and PP2 analogs, which do not inhibit the division of BY-2 cells (Fig. S2 and Table S1). Among the PD-180970 analogs, PD-166326 completely blocked cell division (100%, n = 27), whereas the PD-173955-Analog1 showed only faint effects (13%, n = 63), which were not significant compared to DMSO-treated samples (3%, n = 60, *P* = 0.1) (Fig. S2A and Table S1). Therefore, we chose PD-173955-Analog1 as a negative control for PD-180970.

Among the PP2 analogs, only PP1 inhibited cell division (100%, n = 32), whereas the other analogs had no inhibitory effect on cell division, as represented by PP3 (0%, n = 20, Fig. S2B and Table S1). Therefore, PP3 was selected as the negative control for PP2. We also tested known inhibitors of Bcr-Abl (ponatinib, bosutinib, and bafetinib) and Src kinase (Src inhibitor1) in BY-2 cells, and all exhibited zero or partial inhibition (0–70%, Fig. S2A and Table S1). These results support our hypothesis that the inhibitory targets of PD-180970 and PP2 are plant-specific and independent of the Bcr-Abl and Src kinases found in animals.

PD-180970 and PP2 directly bind to core kinase pockets in animals and act as strong ATP-competitive inhibitors (20, 21, 23–25). Therefore, we hypothesized that these compounds target certain plant kinases and prevent the phosphorylation of particular substrates that are crucial for MT organization and phragmoplast formation (Fig. 2A). These substrates are expected to be abundant in dividing cells, and their phosphorylation levels are reduced in the presence of PD-180970 and PP2. To identify substrates, we performed phosphoproteomics using BY-2 cells (Fig. 2B). The BY–GTRC strain was synchronized at the mitosis (M) phase in the presence of the effective compound (PD-180970 or PP2) or ineffective controls (PD-173955-Analog1 or PP3), and whole cellular proteins were extracted. The phosphopeptides were then purified, and their sequences were determined using a high-sensitivity nanoLC-MS/MS system. In parallel, we generated a BY-2 protein reference database consisting of 50,171 sequences by converting the published transcriptome (RNA-seq) data obtained from non-transgenic BY-2 cells into amino acid sequences (Fig. 2B) (19). There were 14 and 12 proteins identified as the target candidates for PD-180970 and PP2, respectively, based on the criteria that the identified phosphopeptide number was 10 or more in the control cells with ineffective analogs and reduced to less than half in the presence of effective compounds (Datasets S1 and S2, top five candidates are shown in Fig. 2C and D, respectively). Candidate proteins were subjected to subsequent analyses to predict the target events of PD-180970 and PP2.

**Figure 2.**
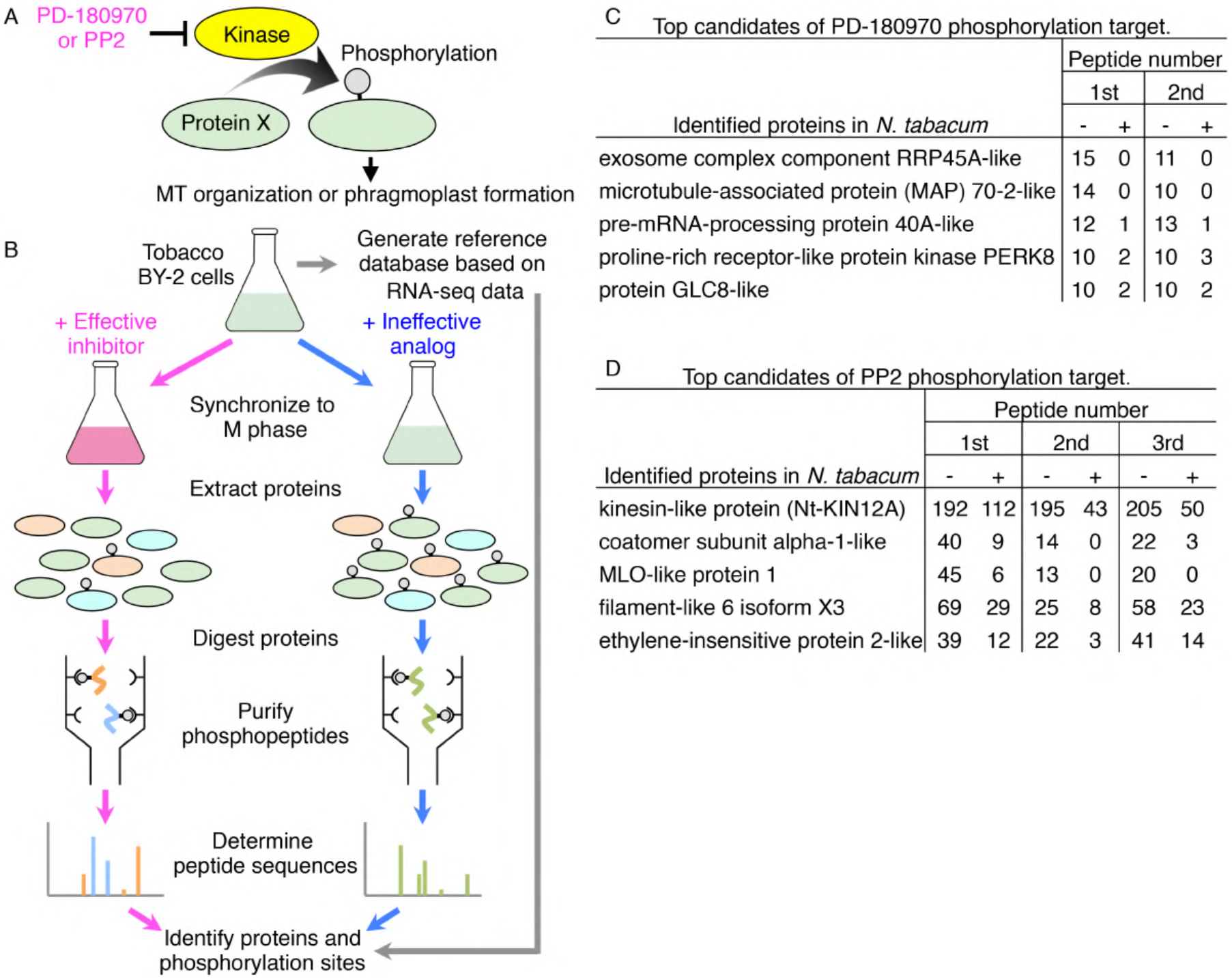
Target identification of PD-180970 and PP2. (A) Hypothetical model of the inhibitory mechanism of PD-180970 and PP2. (B) Schematic procedure of phosphoproteomics to identify the phosphorylation substrate (‘protein X’ in (A)). (C and D) Top 5 candidates of phosphorylation target of PD-180970 (C) and PP2 (D). Peptide number shows the count of identified phosphopeptides in two experiments with effective PD-180970 (+) and ineffective PD-173955-Analog1 (-) (C), and in three experiments with effective PP2 (+) and ineffective PP3 (−) (D). All identified candidates and detailed data are shown in Datasets S1 and S2.

### MAP70 deemed potent candidate of PD-180970 target

Among the identified candidates for PD-180970 targets, RIBONUCLEASE PH45A-like (Nt-RRP45A-like) and MICROTUBULE-ASSOCIATED PROTEIN70-2-like (Nt-MAP70-2-like) showed no phosphopeptides in the presence of PD-180970, suggesting a strong inhibition (Fig. 2C and Dataset S1). The Nt-RRP45A-like candidate exhibited similarity to the RRP45a, CER7, RRP42, and AT1G60080 Arabidopsis proteins, which are predicted to function in RNA-processing/degrading exosomes (26). However, the identified Nt-RRP45A-like phosphorylation sites were not present in Arabidopsis proteins (Fig. S3), which conflicts with the inhibitory effects observed in tobacco BY-2 and Arabidopsis.

In contrast, the three identified phosphorylated serine residues of Nt-MAP70-2-like were present in most Arabidopsis homologs (MAP70-1 to −5) (Fig. S4A). Although the molecular functions of MAP70 remain unclear, MAP70-1 and −5 have reportedly decorated all MT structures (27, 28). Among the three conserved phosphorylation sites, two were located in the essential region for MT association (Fig. S4A) (28). Therefore, we tested the effect of PD-180970 on MT structures using a high-resolution imaging system in BY-2 cells (8). After 30-60 min treatment of with PD-180970 on MT/histone markers, the cortical MT, PPB, spindle, and phragmoplasts were severely disrupted (Fig. 3A). This rapid and general effect supported our hypothesis that PD180970 disrupts MT organization as the primary effect via direct inhibition of MAP70 phosphorylation within the MT-associating motif. An examination of T-DNA insertion mutants of the five Arabidopsis *MAP70* genes showed no detectable defects in root growth (Fig. S4B), implying a high gene redundancy or the presence of additional PD-180970 targets.

**Figure 3.**
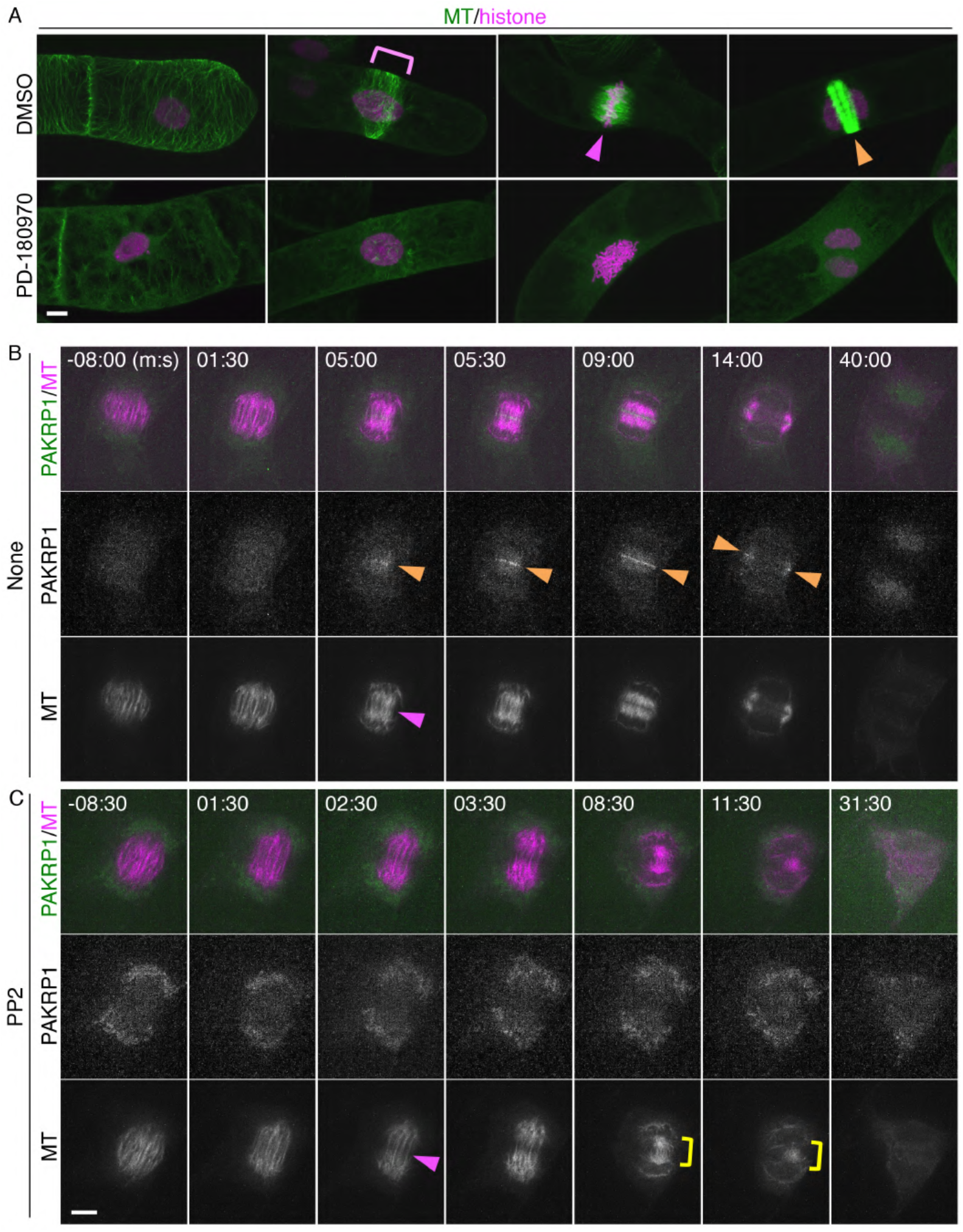
PD-180970 disrupts MT organization and PP2 blocks phragmoplast formation. (A) Confocal images of the BY-2 cells expressing MT/histone marker at 30-60 min after the application of indicated compounds. Magenta rectangle shows PPB. Magenta and orange arrowheads indicate the aligned chromatin at the center region of spindle, and the phragmoplast, respectively. (B and C) Time-lapse observation of the BY-2 cells expressing PAKRP1/MT marker in the presence of no compounds (B) and PP2 (C). Numbers indicate the time (min:sec) from the anaphase onset. Upper, middle, and lower panels show the merged, PAKLP1, and MT images, respectively. Magenta and orange arrowheads indicate the central gap region of spindle and the PAKRP1 localization on phragmoplast, respectively. Yellow rectangle shows the remnant MT bundles connecting two sister chromatids. Scale bars: 10 μm.

### Class II Kinesin-12 deemed potent candidate of PP2 target

Among the identified candidates of PP2 targets, KINESIN-12A (Nt-KIN12A) showed over 100 phosphopeptides in the control cells (Fig. 2D and Dataset S2), with protein abundance and/or high phosphorylation levels in the dividing cells. In Arabidopsis, PHRAGMOPLAST-ASSOCIATED KINESIN-RELATED PROTEIN1 (PAKRP1)/KIN12A and PAKRP1L//KIN12B were the proteins most similar to Nt-KIN12A (Fig. S5). PAKRP1 and PAKRP1L belong to the class II Kinesin-12 family with KIN12F, whose function remains unidentified (29). Kinesin-12 members have MT plus end-directed motility, and PAKRP1 and PAKRP1L are essential for cell plate formation, as *pakrp1 pakrp1l* double mutant disturbs the first postmeiotic cytokinesis owing to disorganization of phragmoplast MTs (2). Twelve phosphorylated residues were identified in Nt-KIN12A, and six were conserved in PAKRP1 or PAKRP1L (Fig. S5). Five conserved residues were found between the putative kinesin motor domain and coiled-coil region (Fig. S5), implying a regulatory element for protein activity and/or interaction (30, 31).

To test whether PP2 disrupts the localization of PAKRP1 and PAKRP1L on phragmoplasts, we generated fluorescent markers and observed co-localization with MT in the presence and absence of the inhibitor using a high-speed time-lapse system of BY-2 cells (PAKRP1/MT, Fig. 3B and Movie S4, and PAKRP1L/MT, Movie S5) (8, 32). In the absence of PP2, both proteins appeared at the central gap region in the remnant spindle, where phragmoplasts emerged (Fig. 3B, and Movies S4 and S5). Both proteins associated with the expanding phragmoplasts then disappeared upon completion. In the presence of PP2, the central gap region was detected, but PAKRP1 and PAKRP1L did not accumulate at this site (Fig. 3C, and Movies S4 and S5). The proper phragmoplasts were not formed throughout the entire process, and the MT bundles were abnormally concentrated in the cell center. These results show that PAKRP1 and PAKRP1L localization was abolished by PP2 as early as the phragmoplast initiation phase. We propose that PP2 blocks phragmoplast formation by inhibiting the phosphorylation of class II Kinesin-12.

### PD-180970 and PP2 do not irreversibly damage viability

To test the utility of PD-180970 and PP2 as cell division inhibitors in physiological experiments, we examined their long-term effects and toxicity. We used Arabidopsis seedlings, which are easily transferred between inhibitor-free and inhibitor-containing media, and whose root meristems show active and regular cell division (Fig. S6) (33). After a 1-day treatment with the compounds, the root tips of the histone marker plants were stained with propidium iodide (PI) to visualize the plasma membrane (histone + PI, Fig. S6A). Compared to DMSO-treated seedlings, oryzalin-treated seedlings had thicker roots consisting of enlarged nuclei (Fig. S6A). Impaired nuclear division caused by oryzalin likely doubled the DNA content and thus increased the nuclear size, as previously reported for the shoot apex (34). The PD-180970-treated seedlings also had enlarged nuclei, whereas the PP2-treated seedlings contained binuclear cells (Fig. S6A). These results showed that their inhibitory effects were consistent with those observed in zygote and BY-2 cells.

We then assessed long-term effects by recording the root length daily in a 4-day treatment (Fig. S6B). Despite their different effects on cells, treatment with PD-180970, PP2, and oryzalin similarly arrested root growth. To test the toxicity of the inhibitors, we compared the root length of seedlings grown in the presence or absence of the inhibitors after a 2-day inhibitor treatment (Fig. S6C). The ratio between root lengths before and after the 5-day incubation was used as a growth indicator. Oryzalin-treated seedlings were unable to grow after inhibitor removal, probably because of the consumption of meristematic cells during treatment. In contrast, the PD-180970- and PP2-treated seedlings successfully restarted their growth (Fig. S6C), indicating that these inhibitors did not cause irreversible damage to the cell viability.

### PD-180970 and PP2 are effective in diverse plant species

To test whether PD-180970 and PP2 could be utilized in other plant species, we analyzed root length following inhibitor treatments in tobacco (*Nicotiana benthamiana*) and cucumber (*Cucumis sativus*). As observed for Arabidopsis (Fig. S6B), all compounds strongly interfered with root growth in both species (Fig. S7A and B). In addition to these angiosperm species, we also tested the moss *Physcomitrium patens*, which diverged from angiosperms at least 500 million years ago (35). Similar to oryzalin, PD-180970 effectively perturbed MT organization, as the filamentous cortical arrays of MT markers were disrupted in the chloronema tip cells of *P. patens* (Fig. 4A) (36). Free GFP-tubulin signals were detected in the cytoplasm, and chloroplast shapes were highlighted in compound-treated cells. This result suggests that PD-180970 affects MT in moss in a manner similar to that observed in Arabidopsis zygotes and tobacco BY-2 cells (Fig. 1H-J and Fig. 3A, respectively).

**Figure 4.**
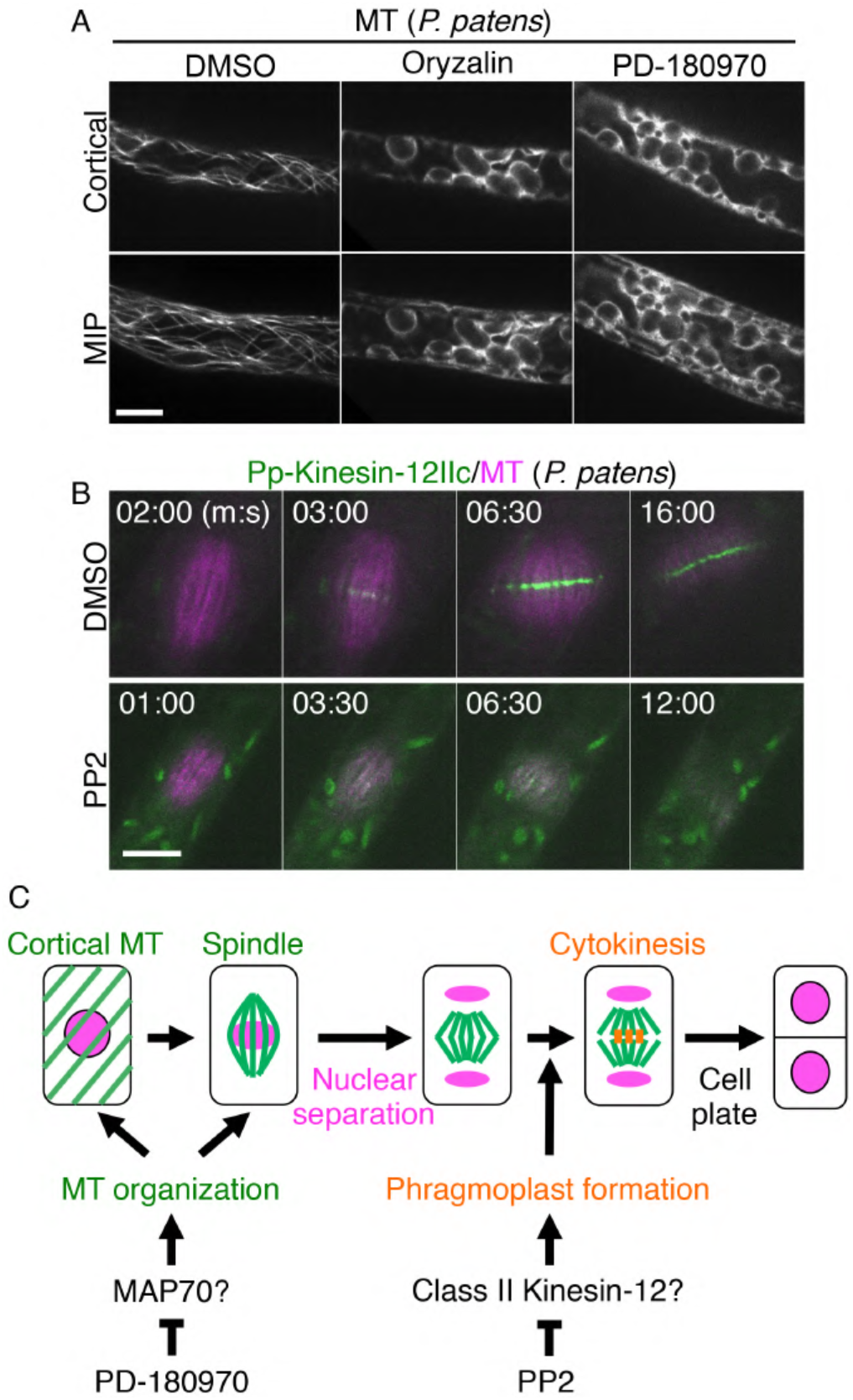
PD-180970 and PP2 are both effective in *P. patens*. (A) Confocal images of *P. patens* chloronema tip cells expressing MT marker at 30 min after the application of indicated compounds. Upper panels show the images of the cortical region, and lower panels display MIP images. (B) Time-lapse observation of *P. patens* chloronema tip cells expressing Pp-Kinesin-12IIc/MT marker in the presence of indicated compounds. Numbers indicate the time (min:sec) from the anaphase onset. (C) Schematic representation of the inhibitory dynamics of PD-180970 and PP2 during plant cell division. Scale bars: 10 μm.

To examine whether PP2 affects class II Kinesin-12 protein in the moss, we observed Pp-Kinesin-12IIc, which localizes at the phragmoplast midzone (Pp-Kinesin-12IIc/MT, Fig. 4B and Movie S6) (37). Pp-Kinesin-12IIc localization was altered by PP2 application, as the strong accumulation at the phragmoplast midzone disappeared (Fig. 4B and Movie S6), consistent with tobacco BY-2 cells (Fig. 3B and C). In the presence of PP2, proper phragmoplast formation was abolished and no cell plate was formed, resulting in binuclear cells and subsequent nuclear fusion (Fig. S7C and Movie S7); both of these results were observed in the BY-2 and Arabidopsis zygotes (Fig. 1G and Fig. S1B). We conclude that PD-180970 and PP2 are effective in blocking MT organization and phragmoplast formation, respectively, and are applicable to diverse plant species (Fig. 4C).

## Discussion

In this study, we identified cell division inhibitors by combining a chemical screening in Arabidopsis zygotes, specification of target events using tobacco BY-2 cells, and validation of their effectiveness in multiple plant species. According to our high-resolution time-lapse imaging and phosphoproteomics, two identified inhibitors, PD-180970 and PP2, were specific to the MT alignment and phragmoplast formation, respectively. In addition, their inhibitory effects were non-lethal and effective in various cell types and plant species. These properties make PD-180970 and PP2 useful tools for future cell division studies, as they provide novel manipulation nodes. We identified MAP70 proteins, as they represent a potential downstream target of PD-180970. MAP70 is an MT-associated factor (38), but its molecular functions are still unknown, except for Arabidopsis MAP70-5, which increases MT length *in vitro* (27). Our phosphoproteomic analysis identified several phosphorylated sites in the conserved MT-binding domain (28), which was consistent with PD-180970 severely disrupting all the tested MT structures (Figs. 1, 3, and 4). These findings suggest that PD-180970 blocks yet-unidentified kinases that phosphorylate MAP70 to disrupt MT organization (Fig. 4C), similar to Aurora kinase, which phosphorylates MAP65-1 to activate its MT-bundling capacity (39). It would be difficult to visualize the PD-180970’s effect on the MT-binding ability of MAP70 because MT alignment itself was destroyed in the presence of PD-180970. More detailed investigations are necessary to reveal the relationship between PD-180970, MAP70, and MT organization. For example, it would be useful to identify the direct binding targets of PD-180970 and to assess whether the phospho-mimic MAP70, whose serines are all replaced with aspartates, becomes tolerant to PD-180970 upon MT disruption. These analyses will also help to detail the molecular function of MAP70 proteins, which are presumably important proteins localized on entire MT structures (27, 28) and are difficult to assess using a genetic approach, owing to the lack of a detectable mutant phenotype (Fig. S4). In addition, we cannot exclude the additional contributions of other candidate proteins whose phosphorylation levels were reduced in the presence of PD-180970 (Fig. 2 and Dataset S1), although we could not find any direct relationship between these proteins and MT organization in the literature.

We predicted class II Kinesin-12 as the top candidate for PP2 through phosphorylation inhibition. In both tobacco BY-2 cells and *P. patens*, PP2 blocked the accumulation of class II Kinesin-12 proteins in the central region of the remnant spindle from the beginning of phragmoplast initiation (Figs. 3 and 4). In Arabidopsis, a double mutant lacking PAKRP1 and PAKRP1L did not form phragmoplast MT during the first postmeiotic cytokinesis (2). These observations were in accordance with the strong inhibition of PP2 on cytokinesis, thereby suggesting class II Kinesin-12 is a potent downstream target of PP2 (Fig. 4C). In addition to class II, the Kinesin-12 family contains three members of class I: PHRAGMOPLAST ORIENTING KINESINS (POK1)/KIN12C, POK2/KIN12D, and KIN12E (29). POK1/POK2 and KIN12E are required for cell plate orientation and spindle assembly, respectively, but they also may localize to the phragmoplast midzone (29, 40, 41). To understand the inhibitory mechanism and specificity of PP2, it would be useful to examine whether PP2 also affects class I members and another class II member, KIN12F, and to identify the direct binding targets of PP2.

In Arabidopsis, PAKRP1 and PAKRP1L interact with the TWO-IN-ONE (TIO)/FUSED kinase (42). Although it remains unclear whether TIO phosphorylates these proteins, the *tio* mutant and *pakrp1 pakrp1l* double mutant failed to form a cell plate during male gametogenesis, suggesting their cooperative function in cytokinesis (2, 43, 44). Therefore, it is important to assess whether TIO is the PP2’s direct binding target to mediate class II Kinesin-12 phosphorylation during phragmoplast formation.

In contrast to the gametogenetic defects of the *pakrp1 pakrp1l* double mutant, individual mutants showed no detectable defects (2, 3). This gene redundancy and mutant lethality of key regulators prohibit our genetic approaches from analyzing the molecular mechanisms of specific cell division events. As a result, even the core regulations, such as how phragmoplasts are initiated and how membrane trafficking toward the cell plate is promoted, are still poorly understood. Both PD-180970 and PP2 showed strong and reversible effects, suggesting that they temporally block related targets simultaneously, thereby circumventing redundancy, lethality, and secondary effects. Therefore, we believe that these compounds are powerful tools for investigating the detailed mechanisms of MT organization and phragmoplast formation.

## Materials and Methods

Detailed materials and methods are described in Supplementary Methods.

### Plant strains and observation

All Arabidopsis lines were observed on a Columbia (Col-0) background. The zygotic histone/PM, MT, and F-actin markers have been described previously (12, 15). These strains were imaged as described previously (14). For live cell imaging of tobacco BY-2 cells, we followed a previously described method (8, 32, 45). The fluorescent markers of *P. patens* and imaging methods have been described previously (36, 37, 46).

### Chemical screening

Young ovules were collected from the siliques of approximately 5 mm and cultivated in 96-well glass bottom plates with individual compounds at 10 μM from the LOPAC® Pfizer library (LO5100; Sigma-Aldrich) and SCREEN-WELL® Kinase Inhibitor library (BML-2832; Enzo Life Science). After incubation for two days in ovule cultivation media (12, 13), the ovules were observed using an inverted fluorescent microscope (IX73; Olympus).

### Phosphoproteomics using synchronized BY-2 culture cells

To perform phosphoproteomics, BY-GTRC cells were synchronized at the DNA replication phase as described previously (47, 48) and then cultured in the presence of the individual compounds for 8-9 h. After confirming that most cells entered the mitosis phase by fluorescent observation, total proteins were extracted and digested with trypsin. Total proteins were digested with trypsin and phosphopeptides were extracted using the IMAC/SIMAC method for nanoLC-MS/MS analysis (49).

## Acknowledgments

We thank Tomomi Yamada, Yumi Kuwabara, Terumi Nishii, and Azusa Utsumi for technical support and Michiko Sasabe, Masaki Ito, Gohta Goshima, Yoshikatsu Sato, Simon Miller, Tsuyoshi Hirota, Shunsuke Oishi, Yasunori Machida, and Ping Kao for helpful discussions. We thank Takeshi Kuroha, Mina Ohtsu, and Michitaka Notaguchi for providing seeds of various plant species and Toshinori Kinoshita and Koji Takahashi for providing the chemical library. Microscopy was supported by the Live Imaging Center at the Institute of Transformative Bio-Molecules (WPI-ITbM) of Nagoya University. This work was supported by Japan Advanced Plant Science Network, Japan Society for the Promotion of Science [Grant-in-Aid for Early-Career Scientists (JP21K15117 for Y.K.), Grant-in-Aid for Research Activity Start-up (19K23723 for M.Y.), Grant-in-Aid for Scientific Research on Innovative Areas (19H04859, 19H05670, and 19H05676 for M.U.; JP16H06465, JP16H06464, JP16K21727 for T.H.; JP16H06280 (Advanced Bioimaging Support); JP20H05358 for D.K.), Grant-in-Aid for Scientific Research (B) (JP19H03243 for M.U. and JP18H02471 for T.M.), Grant-in-Aid for Scientific Research (S) (JP16H06378 for T.M. and M.H.), Grant-in-Aid for Challenging Exploratory Research (JP19K22421 for M.U.)], the Japan Science and Technology Agency [PRESTO programme (JPMJPR18K4 for D.K.), and CREST (JPMJCR2121 for M.U.)], Suntory Rising Stars Encouragement Program in Life Sciences (SunRiSE) for M.U., and Toray Science Foundation to M.U. (20-6102).

## Notes

### Competing Interest Statement

The authors have declared no competing interest.

## References

1. S. Müller, Plant cell division - defining and finding the sweet spot for cell plate insertion. Current opinion in cell biology 60, 9–18 (2019).

2. Y. R. Lee, Y. Li, B. Liu, Two Arabidopsis phragmoplast-associated kinesins play a critical role in cytokinesis during male gametogenesis. The Plant cell 19, 2595–2605 (2007).

3. R. Pan, Y. R. Lee, B. Liu, Localization of two homologous Arabidopsis kinesin-related proteins in the phragmoplast. Planta 220, 156–164 (2004).

4. I. Luptovčiak, G. Komis, T. Takáč, M. Ovečka, J. Šamaj, Katanin: A Sword Cutting Microtubules for Cellular, Developmental, and Physiological Purposes. Frontiers in plant science 8, 1982 (2017).

5. M. Serrano, E. Kombrink, C. Meesters, Considerations for designing chemical screening strategies in plant biology. Frontiers in plant science 6, 131 (2015).

6. J. Y. Ong, J. Z. Torres, Dissecting the mechanisms of cell division. The Journal of biological chemistry 294, 11382–11390 (2019).

7. T. Nagata, Y. Nemoto, S. Hasezawa, Tobacco BY-2 Cell Line as the “HeLa” Cell in the Cell Biology of Higher Plants. 132, 1–30 (1992).

8. T. Murata et al., Mechanism of microtubule array expansion in the cytokinetic phragmoplast. Nature communications 4, 1967 (2013).

9. M. Sasabe et al., Phosphorylation of a mitotic kinesin-like protein and a MAPKKK by cyclin-dependent kinases (CDKs) is involved in the transition to cytokinesis in plants. Proceedings of the National Academy of Sciences of the United States of America 108, 17844–17849 (2011).

10. S. G. Mansfield, L. G. Briarty, Early embryogenesis in Arabidopsis thaliana. II. The developing embryo. Can J Bot 69, 461–476 (1991).

11. M. Ueda et al., Transcriptional integration of paternal and maternal factors in the Arabidopsis zygote. Genes & development 31, 617–627 (2017).

12. K. Gooh et al., Live-cell imaging and optical manipulation of *Arabidopsis* early embryogenesis. Developmental cell 34, 242–251 (2015).

13. D. Kurihara, Y. Kimata, T. Higashiyama, M. Ueda, In Vitro Ovule Cultivation for Live-cell Imaging of Zygote Polarization and Embryo Patterning in Arabidopsis thaliana. Journal of visualized experiments: JoVE 10.3791/55975 (2017).

14. M. Ueda, Y. Kimata, D. Kurihara, Live-Cell Imaging of Zygotic Intracellular Structures and Early Embryo Pattern Formation in Arabidopsis thaliana. Methods in molecular biology (Clifton, N.J.) 2122, 37–47 (2020).

15. Y. Kimata et al., Cytoskeleton dynamics control the first asymmetric cell division in Arabidopsis zygote. Proceedings of the National Academy of Sciences of the United States of America 113, 14157–14162 (2016).

16. Y. Kimata et al., Mitochondrial dynamics and segregation during the asymmetric division of Arabidopsis zygotes. Quantitative Plant Biology 1, e3 (2020).

17. H. Matsumoto, Y. Kimata, T. Higaki, T. Higashiyama, M. Ueda, Dynamic Rearrangement and Directional Migration of Tubular Vacuoles are Required for the Asymmetric Division of the Arabidopsis Zygote. Plant & cell physiology 62, 1280–1289 (2021).

18. A. De Antoni, S. Maffini, S. Knapp, A. Musacchio, S. Santaguida, A small-molecule inhibitor of Haspin alters the kinetochore functions of Aurora B. The Journal of cell biology 199, 269–284 (2012).

19. E. Kozgunova, T. Suzuki, M. Ito, T. Higashiyama, D. Kurihara, Haspin has Multiple Functions in the Plant Cell Division Regulatory Network. Plant & cell physiology 57, 848–861 (2016).

20. J. F. Dorsey, R. Jove, A. J. Kraker, J. Wu, The pyrido[2,3-d]pyrimidine derivative PD180970 inhibits p210Bcr-Abl tyrosine kinase and induces apoptosis of K562 leukemic cells. Cancer Res 60, 3127–3131 (2000).

21. J. H. Hanke et al., Discovery of a novel, potent, and Src family-selective tyrosine kinase inhibitor. Study of Lck-and FynT-dependent T cell activation. The Journal of biological chemistry 271, 695–701 (1996).

22. D. Kurihara, S. Matsunaga, S. Uchiyama, K. Fukui, Live cell imaging reveals plant aurora kinase has dual roles during mitosis. Plant & cell physiology 49, 1256–1261 (2008).

23. P. La Rosee, A. S. Corbin, E. P. Stoffregen, M. W. Deininger, B. J. Druker, Activity of the Bcr-Abl kinase inhibitor PD180970 against clinically relevant Bcr-Abl isoforms that cause resistance to imatinib mesylate (Gleevec, STI571). Cancer Res 62, 7149–7153 (2002).

24. N. Vajpai et al., Solution conformations and dynamics of ABL kinase-inhibitor complexes determined by NMR substantiate the different binding modes of imatinib/nilotinib and dasatinib. The Journal of biological chemistry 283, 18292–18302 (2008).

25. X. Zhu et al., Structural analysis of the lymphocyte-specific kinase Lck in complex with non-selective and Src family selective kinase inhibitors. Structure (London, England: 1993) 7, 651–661 (1999).

26. T. S. Hooker, P. Lam, H. Zheng, L. Kunst, A core subunit of the RNA-processing/degrading exosome specifically influences cuticular wax biosynthesis in Arabidopsis. The Plant cell 19, 904–913 (2007).

27. A. V. Korolev, H. Buschmann, J. H. Doonan, C. W. Lloyd, AtMAP70-5, a divergent member of the MAP70 family of microtubule-associated proteins, is required for anisotropic cell growth in Arabidopsis. Journal of cell science 120, 2241–2247 (2007).

28. A. V. Korolev, J. Chan, M. J. Naldrett, J. H. Doonan, C. W. Lloyd, Identification of a novel family of 70 kDa microtubule-associated proteins in Arabidopsis cells. The Plant journal: for cell and molecular biology 42, 547–555 (2005).

29. S. Müller, P. Livanos, Plant Kinesin-12: Localization Heterogeneity and Functional Implications. International journal of molecular sciences 20 (2019).

30. S. C. Ems-McClung, C. E. Walczak, Kinesin-13s in mitosis: Key players in the spatial and temporal organization of spindle microtubules. Seminars in cell & developmental biology 21, 276–282 (2010).

31. B. J. Mann, P. Wadsworth, Kinesin-5 Regulation and Function in Mitosis. Trends in cell biology 29, 66–79 (2019).

32. T. Murata, T. I. Baskin, Imaging the mitotic spindle by spinning disk microscopy in tobacco suspension cultured cells. Methods in molecular biology (Clifton, N.J.) 1136, 47–55 (2014).

33. P. N. Benfey, J. W. Schiefelbein, Getting to the root of plant development: the genetics of Arabidopsis root formation. Trends in Genetics 10, 84–88 (1994).

34. O. Grandjean et al., In vivo analysis of cell division, cell growth, and differentiation at the shoot apical meristem in Arabidopsis. The Plant cell 16, 74–87 (2004).

35. J. L. Morris et al., The timescale of early land plant evolution. Proceedings of the National Academy of Sciences of the United States of America 115, E2274–e2283 (2018).

36. E. Kozgunova, G. Goshima, A versatile microfluidic device for highly inclined thin illumination microscopy in the moss Physcomitrella patens. Scientific reports 9, 15182 (2019).

37. T. Miki, H. Naito, M. Nishina, G. Goshima, Endogenous localizome identifies 43 mitotic kinesins in a plant cell. Proceedings of the National Academy of Sciences of the United States of America 111, E1053–1061 (2014).

38. L. C. Morejohn, T. E. Bureau, J. Mole-Bajer, A. S. Bajer, D. E. Fosket, Oryzalin, a dinitroaniline herbicide, binds to plant tubulin and inhibits microtubule polymerization in vitro. Planta 172, 252–264 (1987).

39. J. Boruc et al., Phosphorylation of MAP65-1 by Arabidopsis Aurora Kinases Is Required for Efficient Cell Cycle Progression. Plant physiology 173, 582–599 (2017).

40. A. Herrmann et al., Dual localized kinesin-12 POK2 plays multiple roles during cell division and interacts with MAP65-3. EMBO reports 19 (2018).

41. A. Herrmann et al., KINESIN-12E regulates metaphase spindle flux and helps control spindle size in Arabidopsis. The Plant cell 33, 27–43 (2021).

42. S. A. Oh et al., Arabidopsis Fused kinase and the Kinesin-12 subfamily constitute a signalling module required for phragmoplast expansion. The Plant journal: for cell and molecular biology 72, 308–319 (2012).

43. S. A. Oh, V. Bourdon, H. G. Dickinson, D. Twell, S. K. Park, Arabidopsis Fused kinase TWO-IN-ONE dominantly inhibits male meiotic cytokinesis. Plant Reprod 27, 7–17 (2014).

44. S. A. Oh et al., A divergent cellular role for the FUSED kinase family in the plant-specific cytokinetic phragmoplast. Current biology: CB 15, 2107–2111 (2005).

45. M. Nambo et al., Combination of Synthetic Chemistry and Live-Cell Imaging Identified a Rapid Cell Division Inhibitor in Tobacco and Arabidopsis thaliana. Plant & cell physiology 57, 2255–2268 (2016).

46. M. Yamada, T. Miki, G. Goshima, Imaging Mitosis in the Moss Physcomitrella patens. Methods in molecular biology (Clifton, N.J.) 1413, 263–282 (2016).

47. T. Nagata, F. Kumagai, Plant cell biology through the window of the highly synchronized tobacco BY-2 cell line. Methods in cell science: an official journal of the Society for In Vitro Biology 21, 123–127 (1999).

48. F. Kumagai-Sano, T. Hayashi, T. Sano, S. Hasezawa, Cell cycle synchronization of tobacco BY-2 cells. Nat Protoc 1, 2621–2627 (2006).

49. Y. Ohkubo, K. Kuwata, Y. Matsubayashi, A type 2C protein phosphatase activates high-affinity nitrate uptake by dephosphorylating nRT2.1. Nature plants 10.1038/s41477-021-00870-9 (2021).

