## Supplementary material for "Chemical screen of Arabidopsis zygote and proteomics in tobacco BY-2 cells identify general plant cell division inhibitors": Figure S

### **Supplementary Methods**

### **Growth conditions and plant strains**

Arabidopsis and *N. benthamiana* were grown in Petri dishes containing a 1.5% agar medium and 1/2 Murashige and Skoog (MS) medium or on soil at 18–22 °C under continuous light or long-day conditions (16 h light/8 h dark). Tobacco BY-2 cells and *P. patens* strains were cultured as described previously (1, 2). The 'Chinese long' strain was used as *C. sativus* and grown in Petri dishes containing 0.1% PLANT PRESERVATIVE MIXTURE (PPM; nacalai tesque) as antibacterial reagent under continuous light at 18–22 °C.

All Arabidopsis lines were placed in a Columbia (Col-0) background. Mutants of *map70-1* (SALK_013866), *map70-2* (SALK_060997), *map70-3* (SALK_203128), *map70-4* (SALK_069552), and *map70-5* (SALK_106968) were identified in the SALK institute.

### **Compounds**

The following compounds were used; oryzalin (36182, Sigma-Aldrich), 5-Iodotubercidin (5-ITu) (I100, Sigma-Aldrich), PD-180970 (PZ0142, Sigma-Aldrich), PD-166326 (9000988, Cayman Chemical), PD-173955-Analog1 (SYN-1062, SYNkinase), Ponatinib (CS-0204, CHEMSCENE), Bosutinib (PZ0192, Sigma-Aldrich), Bafetinib (A10119, AdooQ Bioscience), PP2 (P0042, Sigma-Aldrich), PP1 (BML-EI275, Enzo Life Science), 4-aminopyrazolo[3,4-d]pyrimidine (PP2-Analog1) (A1041, Tokyo Chemical Industry), 1-tert-butyl-1H-pyrazolo[3,4-d]pyrimidine-4-amine (PP2-Analog2) (GF-0723, Key Organics), PP3 (A2737, Tokyo Chemical Industry), and Src inhibitor1 (sc-204303, Santa Cruz Biotechnology) (3-6).

Each compound was dissolved in DMSO and used at a final concentration of 10 µM in 0.1% DMSO, except for the dosage assay and oryzalin, which was used at 1 µM. Oryzalin (10 µM) was used in the *P. patens* assay.

### **Fluorescent markers and microscopy for Arabidopsis**

In the histone/PM marker for Arabidopsis zygotes and embryos, green fluorescent protein fused to histone H2B and red fluorescent protein fused to LOW-TEMPERATURE-INDUCED6b were expressed under a zygote/embryo-specific promoter of *WUSCHEL RELATED HOMEOBOX2* gene (WOX2p::H2B–GFP and WOX2p::tdTomato–LTI6b) (7). Live-cell imaging of this histone/PM marker was performed using an inverted confocal microscope system (CV1000; Yokogawa Electric) as described previously (8).

MT and F-actin were observed in Arabidopsis zygotes using EC1p::Clover–TUA6 (9) and EC1p::Lifeact–Venus (10), respectively. Live-cell imaging of these markers was performed using a two-photon excitation microscope (LSM780-DUO-NLO; Zeiss) as previously described (11, 12).

For the observation of root meristem in Arabidopsis, the histone marker RPS5Ap::H2B–GFP (13) was stained with 10 µg/ml propidium iodide (PI; Sigma) to visualize the plasma membrane. The samples were observed using an LSM780-DUO-NLO (Zeiss), as described previously (14).

### **Fluorescent markers and microscopy for tobacco BY-2 cells**

As the MT/centromere marker for tobacco BY-2 cell culture cells, we used the BY–GTRC strain, which labeled MT and centromeric histone by expressing GFP–fused α-tubulin and RFP (red fluorescent protein)–labeled CenH3 (centromeric histone H3) (35S::GFP–α-tubulin and

35S::RFP–CenH3) (1). This marker was observed using a confocal microscope system (CV1000; Yokogawa Electric) as described previously (8).
As MT/histone markers for high-resolution imaging of BY-2 cells, we used NOSp::mCitrine–TUB8 and H2Bp::H2B–mCherry. These markers were tandemly cloned into the pCAMBIA1300 binary vector via *P. patens* intergenic (PIG) 1 L region (15). For NOSp::mCitrine–TUB8, a 0.18-kb NOS promoter was fused to yellow fluorescent protein mCitrine, NtTUB8 (LOC107786877) cDNA, and a 0.25-kb AtHSP18.2 (AT5G59720) terminator. For H2Bp::H2B–mCherry, red fluorescent protein mCherry was inserted before the stop codon of genomic sequence of *N. tabacum* histone H2B spanning from 1.9-kb 5'-UTR to 0.5-kb 3'-UTR. Images were acquired as z-stacks with 0.8 µm intervals using a confocal microscope system (FV3000; Olympus) equipped with two GaASP detectors and a water immersion 60× objective lens (NA 1.2). The emission signal of mCitrine was detected between 500 and 550 nm with a 488 nm excitation, and that of mCherry was detected between 600 and 680 nm with a 561 nm excitation. Control DMSO or PD-180970 were applied between 30 and 60 min before observation.
To observe the MT array and Kinesin-12 localization in BY-2 cells, NOSp::mScarlet-i–TUB8 was combined with PAKRP1 and PAKRP1L markers (PAKRP1/MT and PAKRP1L/MT). In NOSp::mScarlet-i–TUB8, the NOS promoter was fused to the red fluorescent protein mScarlet-i, NtTUB8 cDNA, and the 0.47-kb NtEF-1-alpha terminator in a pRI910 binary vector. The PAKRP1 and PAKRP1L markers used were RPS5Ap::Clover–PAKRP1 (MU2384) and RPS5Ap::Clover–PAKRP1L (MU2403), respectively. Among these markers, a 1.7-kb RIBOSOMAL PROTEIN SUBUNIT 5A (RPS5A) promoter (16) was fused to the green fluorescent protein Clover, the full-length coding region of PAKRP1 (AT4G14150) or PAKRP1 (AT3G23670), and the NOPALINE SYNTHASE (NOS) terminator in a pMDC99 binary vector (17). Images were acquired as previously described using an inverted microscope (IX81; Olympus) equipped with a spinning-disk unit (CSU21; Yokogawa) and a water immersion 60× objective lens (NA 1.2), with an additional optical unit (W-view Gemini; Hamamatsu) used for simultaneous image acquisition of green and red channels (18). The high-speed time-lapse images of the PAKRP1/MT and PAKRP1L/MT markers were acquired every 30 s with bandpass filters (Semrock FF01-520/60-25 and FF01-609/54-25 for mClover and mScarlet-i, respectively), and the images were expanded by an additional 1.5× extension lens in front of the camera. The compounds were applied during live-cell imaging as described previously (19).

**Fluorescent markers and microscopy for *P. patens***
*P. patens* chloronema was observed using the MT marker (PpGCP4p::GFP–tubulin) (20), the dual-color marker of MT and nucleus (PpGCP4p::GFP–tubulin and 7113p::histone H2B–mRFP) (20), or Pp-Kinesin-12IIc and MT (Pp-Kinesin-12IIc–Citrine and PpACTp::mCherry–tubulin) (21). For the PD-180970 experiment, chloronema tissues expressing MT marker cultured on a cellophane-laid BCDAT plate for 6 days were sonicated in a BCD liquid medium containing 10 µM oryzalin, 10 µM PD-180970, or 0.5% DMSO, followed by incubation for 30 min. The tissues were introduced into microfluidic devices and immediately observed (20). The images were acquired with an inverted microscope (Ti; Nikon, 100× 1.45 NA lens) equipped with a spinning-disk, confocal unit (CSU-X1; Yokogawa), 488 nm and 561 nm laser lines (LDSYS-488/561-50-YHQSP3, Pneum), and an electron-multiplying charge-coupled device camera (ImagEM ; Hamamatsu) at 2.5 µm z-intervals. The microscope was controlled using NIS elements.
For PP2 experiments, chloronema tissues were cultured in 6-well glass-bottom dishes or 35 mm dishes in a BCD agarose medium for 5-7 days (2). Water containing 10 µM PP2 or 0.5% DMSO was directly applied to the dishes, and the mosses were incubated for 30 min before observation. High-resolution live-cell imaging of the Pp-Kinesin-12IIc/MT marker was performed using the same microscope described above. Long-term imaging of MT/histone markers was performed with a wide-field microscope (TE2000; Nikon, 10x 0.45 NA lens) equipped with a CMOS camera (ZYLA-4.2P-USB3; Andor) and a Nikon Inte4nsilight Epi-fluorescence illuminator, which was controlled by iQ software.

**Phosphoproteomics of tobacco BY-2 cells**
For phosphoproteomics, BY-GTRC cells at 7 days after transfer to fresh medium were synchronized at the DNA replication stage (synthesis (S) phase), as described previously (22, 23).

The cells were cultured in the presence of PD-180970, PD-173955-Analog1, PP2, or PP3 for 8-9 h. After confirming that most cells started mitosis using an upright microscope (AxioImager A2; Zeiss), total proteins were extracted using cell lysis buffer [50 mM Tris-HCl (pH8.0), 150 mM NaCl, 1% (v/v) Triton X-100, 25 µM MG-132, and cOmplete Mini Protease Inhibitor Cocktail (Roche)]. After the extraction of crude proteins and trypsin digestion, peptides were purified using an immobilized metal ion affinity chromatography (IMAC) column or a sequential enrichment of immobilized metal affinity chromatography (SIMAC) column, both of which specifically absorb phosphopeptides. Their amino acid sequences were determined using a high-sensitivity nanoLC-MS/MS system, as previously described (24).
The protein sequences of BY-2 cells were predicted based on the transcriptome (RNA-seq) data, which have been previously reported (25). Each transcript data were converted to an amino acid sequence, and a total of 50,171 protein sequences (coded from NtBYT000000.000 to NtBYT078147.000) were used as a reference database to map the identified phosphopeptides by MASCOT search to determine the corresponding proteins. Identified proteins and their homologous proteins were aligned using the Crustal Omega software (https://www.ebi.ac.uk/Tools/msa/clustalo/). Predictions of the MT-binding region in MAP70 proteins have been previously reported (26). Kinesin motor domains and coiled-coil domains in PAKRP1 and PAKRP1L were predicted using UniProt (https://www.uniprot.org).

**References for Supplementary Methods**


1. D. Kurihara, S. Matsunaga, S. Uchiyama, K. Fukui, Live cell imaging reveals plant aurora kinase has dual roles during mitosis. *Plant & cell physiology* **49**, 1256-1261 (2008).
2. M. Yamada, T. Miki, G. Goshima, Imaging Mitosis in the Moss Physcomitrella patens. *Methods in molecular biology (Clifton, N.J.)* **1413**, 263-282 (2016).
3. J. M. Hum, R. N. Day, J. P. Bidwell, Y. Wang, F. M. Pavalko, Mechanical loading in osteocytes induces formation of a Src/Pyk2/MBD2 complex that suppresses anabolic gene expression. *PloS one* **9**, e97942 (2014).
4. R. Karni, Y. Dor, E. Keshet, O. Meyuhas, A. Levitzki, Activated pp60c-Src leads to elevated hypoxia-inducible factor (HIF)-1alpha expression under normoxia. *The Journal of biological chemistry* **277**, 42919-42925 (2002).
5. F. Rossari, F. Minutolo, E. Orciuolo, Past, present, and future of Bcr-Abl inhibitors: from chemical development to clinical efficacy. *Journal of hematology & oncology* **11**, 84 (2018).
6. D. Wisniewski *et al.*, Characterization of potent inhibitors of the Bcr-Abl and the c-kit receptor tyrosine kinases. *Cancer Res* **62**, 4244-4255 (2002).
7. K. Gooh *et al.*, Live-cell imaging and optical manipulation of *Arabidopsis* early embryogenesis. *Developmental cell* **34**, 242-251 (2015).
8. M. Nambo *et al.*, Combination of Synthetic Chemistry and Live-Cell Imaging Identified a Rapid Cell Division Inhibitor in Tobacco and Arabidopsis thaliana. *Plant & cell physiology* **57**, 2255-2268 (2016).
9. Y. Kimata *et al.*, Cytoskeleton dynamics control the first asymmetric cell division in Arabidopsis zygote. *Proceedings of the National Academy of Sciences of the United States of America* **113**, 14157-14162 (2016).
10. T. Kawashima *et al.*, Dynamic F-actin movement is essential for fertilization in Arabidopsis thaliana. *eLife* **3** (2014).
11. D. Kurihara, Y. Kimata, T. Higashiyama, M. Ueda, In Vitro Ovule Cultivation for Live-cell Imaging of Zygote Polarization and Embryo Patterning in Arabidopsis thaliana. *Journal of visualized experiments : JoVE* 10.3791/55975 (2017).
12. M. Ueda, Y. Kimata, D. Kurihara, Live-Cell Imaging of Zygotic Intracellular Structures and Early Embryo Pattern Formation in Arabidopsis thaliana. *Methods in molecular biology (Clifton, N.J.)* **2122**, 37-47 (2020).
13. D. Maruyama *et al.*, Independent control by each female gamete prevents the attraction of multiple pollen tubes. *Developmental cell* **25**, 317-323 (2013).

14. Y. Kimata *et al.*, Polar vacuolar distribution is essential for accurate asymmetric division of Arabidopsis zygotes. *Proceedings of the National Academy of Sciences of the United States of America* **116**, 2338-2343 (2019).
15. Y. Okano *et al.*, A polycomb repressive complex 2 gene regulates apogamy and gives evolutionary insights into early land plant evolution. *Proceedings of the National Academy of Sciences of the United States of America* **106**, 16321-16326 (2009).
16. S. Adachi *et al.*, Programmed induction of endoreduplication by DNA double-strand breaks in Arabidopsis. *Proceedings of the National Academy of Sciences of the United States of America* **108**, 10004-10009 (2011).
17. M. D. Curtis, U. Grossniklaus, A gateway cloning vector set for high-throughput functional analysis of genes in planta. *Plant physiology* **133**, 462-469 (2003).
18. T. Murata *et al.*, Mechanism of microtubule array expansion in the cytokinetic phragmoplast. *Nature communications* **4**, 1967 (2013).
19. T. Murata, T. I. Baskin, Imaging the mitotic spindle by spinning disk microscopy in tobacco suspension cultured cells. *Methods in molecular biology (Clifton, N.J.)* **1136**, 47-55 (2014).
20. E. Kozgunova, G. Goshima, A versatile microfluidic device for highly inclined thin illumination microscopy in the moss Physcomitrella patens. *Scientific reports* **9**, 15182 (2019).
21. T. Miki, H. Naito, M. Nishina, G. Goshima, Endogenous localizome identifies 43 mitotic kinesins in a plant cell. *Proceedings of the National Academy of Sciences of the United States of America* **111**, E1053-1061 (2014).
22. T. Nagata, F. Kumagai, Plant cell biology through the window of the highly synchronized tobacco BY-2 cell line. *Methods in cell science : an official journal of the Society for In Vitro Biology* **21**, 123-127 (1999).
23. F. Kumagai-Sano, T. Hayashi, T. Sano, S. Hasezawa, Cell cycle synchronization of tobacco BY-2 cells. *Nat Protoc* **1**, 2621-2627 (2006).
24. Y. Ohkubo, K. Kuwata, Y. Matsubayashi, A type 2C protein phosphatase activates high-affinity nitrate uptake by dephosphorylating NRT2.1. *Nature plants* 10.1038/s41477-021-00870-9 (2021).
25. E. Kozgunova, T. Suzuki, M. Ito, T. Higashiyama, D. Kurihara, Haspin has Multiple Functions in the Plant Cell Division Regulatory Network. *Plant & cell physiology* **57**, 848-861 (2016).
26. A. V. Korolev, J. Chan, M. J. Naldrett, J. H. Doonan, C. W. Lloyd, Identification of a novel family of 70 kDa microtubule-associated proteins in Arabidopsis cells. *The Plant journal : for cell and molecular biology* **42**, 547-555 (2005).

**Supplementary Figures**

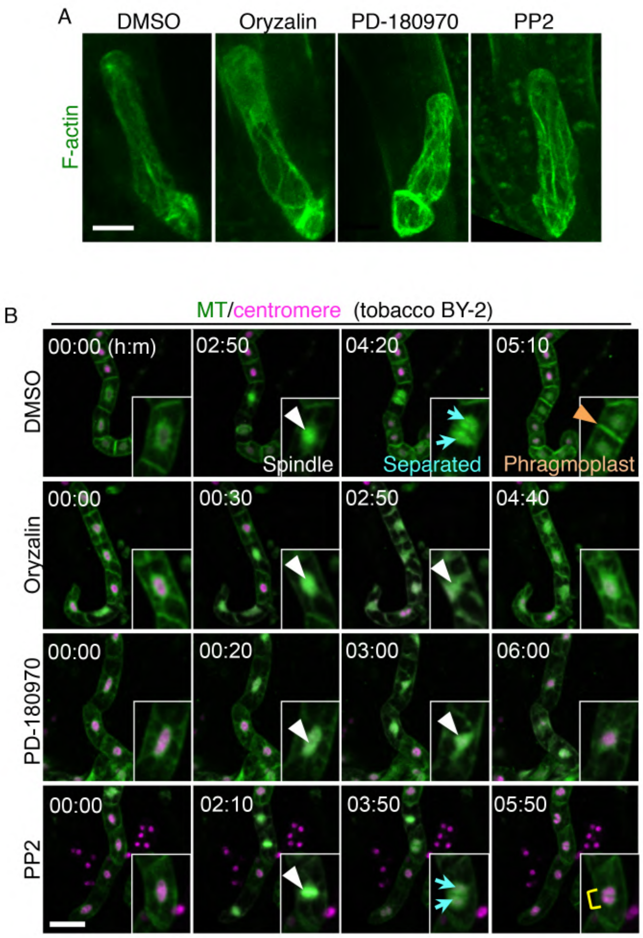


**Fig. S1.** The effect of PD-180970 and PP2 on F-actin in Arabidopsis zygote and tobacco BY-2 cells.

(A) 2PEM images of the Arabidopsis zygotes, expressing F-actin marker at 1 hour after the application of indicated compounds. MIP images are shown. (B) Time-lapse observation of tobacco BY-GTRC cell strain in the presence of the indicated compounds. MIP images are shown. Numbers indicate the time (h:min) from the first frame, and insets show the enlarged images of perinuclear regions. White and orange arrowheads indicate the spindle and phragmoplast, respectively. Cyan arrows and yellow rectangle show separated and two associated nuclei, respectively. Scale bars: 10 μm (A) and 100 μm (B).

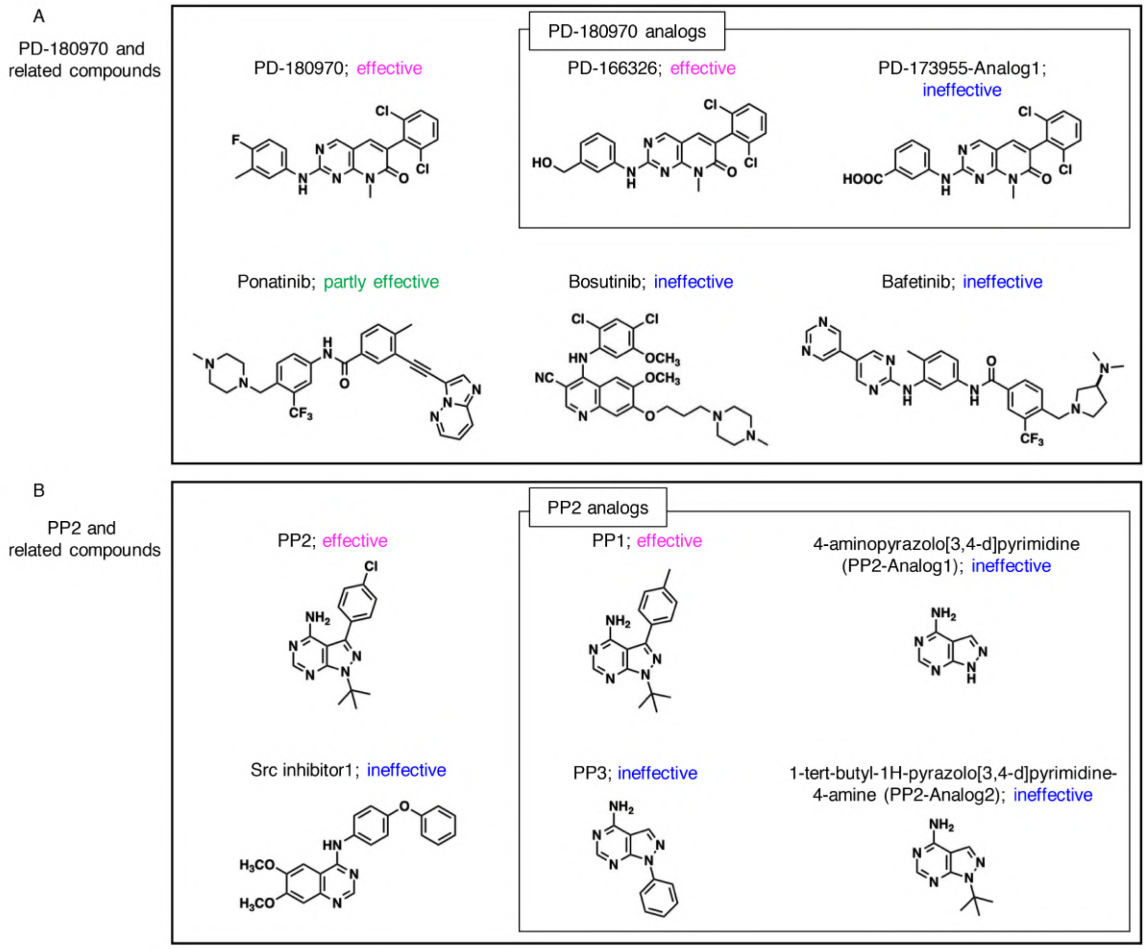


**Fig. S2.** Relative compounds of PD-180970 and PP2.

(A and B) Structure of related compounds to PD-180970 (A) and PP2 (B). Antiproliferative effect of each compound is shown based on the analysis of tobacco BY-2 cells.

```
Nt-RRP45A-like    -----MEQRLANTWR----MTVNEKKFIESALESELRIDGRRPFDYRRLTIKFG---RED    48
RRP45A            -----MEGRLNNMWR----LTVNESKFVESALQSELRVDGRGLYDYRKLTIKFG---KEY    48
CER7              -----MEGRLANMWR----LTVNESKFVETALQSELRVDGRGLYDYRKLTIKFG---KEY    48
RRP42             -------------MG----LSLGEQHFIKGGIAQDLRTDGRKRLTYRPIYVETGVIPQAN    43
AT1G60080         MGIPDAAQDLSTEMEVDAFRRIFPLRFFERHLSESLRPDGRQLGKARDTIVNLGLVSTAD    60

Nt-RRP45A-like    GSSEVQLGQTHVMGFVTSQLVQPYRDRPNEGTLSVFTEFSPMADPSFEAGRPSEYAVELG    108
RRP45A            GSSQVQLGQTHVMAFVTAQLVQPYKDRPSEGSFSIFTEFSPMADPSFEPGHPGESAVELG    108
CER7              GSSEVQLGQTHVMGFVTAQLVQPYKDRPNEGSLSIFTEFSPMADPSFEPGRPGESAVELG    108
RRP42             GSARVRIGGTDVIASVKAEIGRPSSLQPDKGKVAVFIDCSPTAEPTFGGRGGEELSSELA    103
AT1G60080         GSALAKIGSTTMLAAIRMEVMTPSTDSPDEGCIAIEFHMPPICSPTVRPGRPAEAAPVIS    120

Nt-RRP45A-like    RIIDRGLR-----ESRAVDTESLCVVAGKWVWSIRIDLHILDNGGNLVDAANVAALAALL    163
RRP45A            RIIDRALR-----ESRAVDTESLCVLAGKLVWSVRIDLHILDNGGNLVDAANVAALAALM    163
CER7              RIIDRGLR-----ESRAVDTESLCVLAGKMVWSVRIDLHILDNGGNLVDAANIAALAALM    163
RRP42             LALQRCLLGGKSGAGAGINLSSLLIKEGKVCWDLYIDGLVISSDGNLLDALGAAIKAALT    163
AT1G60080         KRLSDTIL-----SSGMIDLKELCLVSGKAAWMGYLDIYCLDADGALFDAALLAAVAAFS    175

Nt-RRP45A-like    TFRRPECTLGGDDRQEVILHPPEVPFLA-----YFPQFFFKPILQKKMFXVTFAFIGD-E    217
RRP45A            TFRRPDCTVGGDNSQDVIIHPPEEREPL-----P-------LIIHHLPIAFTFGFFNKGS    211
CER7              TFRRPDCTVGGENGQEVIIHPLEEREPL-----P-------LIIHHLPIAFTFGFFNKGN    211
RRP42             NTAIPKVNVSAEVADD------EQPEID-----ISDEEYLQFDTSSVPV--IVTLTKVGT    210
AT1G60080         NLQIPIVALNDNGRIVAVTGEKDQDNALITEKEAVNKEKRKLTLKNIPF–SLTCILHKN     233

Nt-RRP45A-like    NLVIDPTHHEEAVMGGRMSVTLNTNGEVCAFQKPGGQGTMQ-SVVMQCLRIASVKAGDIT    276
RRP45A            ILVMDPTYVEEAVMCGRMTVTVNANGDICAIQKPGEEGVNQ-SVILHCLRLASSRASATT    270
CER7              IVVMDPTYVEEAVMCGRMTVTVNANGDICAIQKPGEEGVNQ-SVILHCLRLASSRAAATT    270
RRP42             HYIVDATAEEESQMSSAVSISVNRTGHICGLTKRSGSGLDP-SVILDMISVAKHVTETLM    269
AT1G60080         YILADPTTEEESIMDTLVTVVLDSSDQMVSFYKSGGAALAYSSAIKSCVELARKRAKELK    293

Nt-RRP45A-like    SKIKTAVESY-NTERALKMVKRHTNINAVDVNEPDQKAKQILEGKKL--KSEECSVSQSD    333
RRP45A            KIIRDAVEAY-NRERSSQKVKRHHTLAKSEVLGPIVVV----------------------    307
CER7              KIIREEVEAY-NCERSLQKVKRHPTLAKSEVSGPTVAVKEEHRKSSDQERAAEISREHVE    329
RRP42             SKLDSEISAAEACEDES-------------------------------------------    286
AT1G60080         QILGEMDID---------------------------------------------------    302

Nt-RRP45A-like    DMDVEQGRV-KKRSDKNQRFTGGPSSWDPYSEGVNTDELKATLASRGSAAVPMKLDNLGD    392
RRP45A            ------------------------------------------------------------    307
CER7              RLKLSTEEVRSSKEEEAANFKGGPSNWDPYSEAMDVDSLKVSLASRGDPVTKSSSTKKMN    389
RRP42             ------------------------------------------------------------    286
AT1G60080         ------------------------------------------------------------    302
                         ▼    ▼  ▼
Nt-RRP45A-like    DITDERKSDEPLSDVSPVSSLTESAGKE–MATSREKTLQDAVKPKNRRRKKKSSNSAV      449
RRP45A            ----------------------------------------------------------      307
CER7              GSGNAQKVGV----EISVEEVTGELGKKDTKHKDGEMTLKDAVKPKKKRKNKS-----      438
RRP42             ----------------------------------------------------------      286
AT1G60080         ----------------------------------------------------------      302
```

**Fig. S3.** Alignment of the amino acid sequence of Nt-RRP45A-like with its presumptive homologs in Arabidopsis.

The compared proteins were Nt-RRP45A-like (GenBank ID: XP_016492014.1) from *N. tabacum* and RRP45A (AT3G12990), CER7 (AT3G60500), RRP42 (AT3G07750), and AT1G60080 from Arabidopsis. The similarity (identity) of the full-length amino acid sequence of NtRRP45A-like with each Arabidopsis protein was 60%, 56%, 24%, and 23%, respectively. The identified phosphorylated residues are shown in red with the arrowheads.

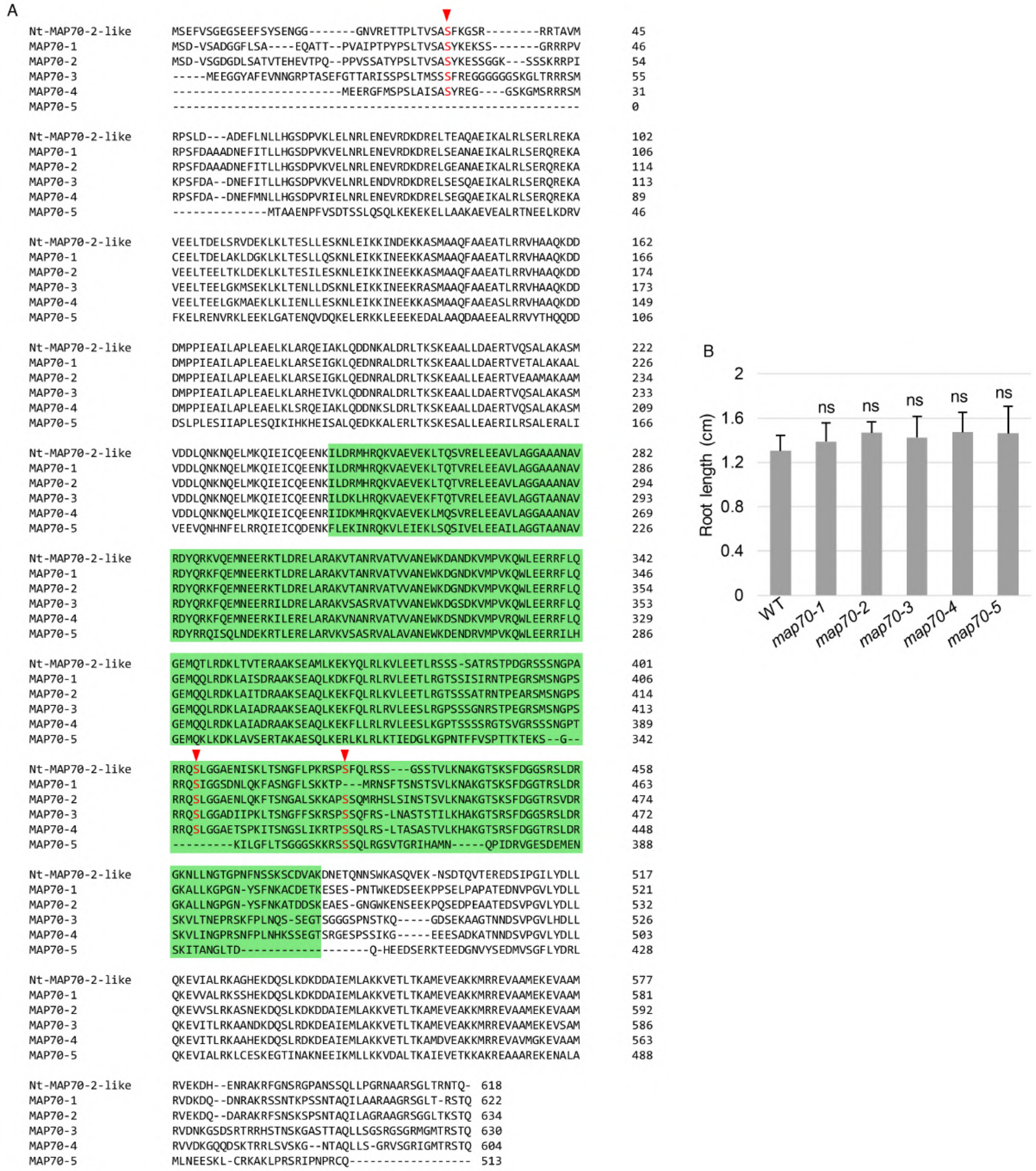


**Fig. S4.** Alignment of the amino acid sequence of Nt-MAP70-2-like with Arabidopsis MAP70-1 to -5, and the phenotype of Arabidopsis mutants.

(A) Compared proteins are Nt-MAP70-2-like (Genbank ID: XP_016475052.1) from *N. tabacum*, and all five MAP70 proteins from Arabidopsis (MAP70-1 (AT1G68060), MAP70-2 (AT1G24764), MAP70-3 (AT2G01750), MAP70-4 (AT1G14840), and MAP70-5 (AT4G17220)). The similarity (identity) of full-length amino acid sequence of Nt-MAP70-2-like to each Arabidopsis protein is 66%, 66%, 64%, 64%, and 40%, respectively. Identified phosphorylated residues are shown in red characters with arrowheads. Putative MT-binding region is green-colored. (B) Root length of 5-day-old seedlings in wild type (WT) and single mutants of *MAP70-1* to *-5*. Error bars represent SD (n = 10). Significant difference was determined by Dunnett's test. Ns, not significant.

```
Nt-KIN12A    ------MSENRFLGNISASSIRNLLPKSVSTKKKLNSSRFKHKMNSENVAPIDPNIQITD   54
PAKRP1       MKKHFTLPRNAILRDGGEP-------HSPNPSISKSKPPRKLRSAKENAPPLDRNTSTPD   53
PAKRP1L      -MKHFMMPRNAILRDIGES-------QSPNPSLTKSKSQRKIKSSKENAPPPDLNSLIPD   52

Nt-KIN12A    PPLLPTSSILKKPVVDTDYTSED------------LTRSEALEQTLEAPDSPVKVVVRIR   102
PAKRP1       HRSM--RMKNPLP--PRPPPSNP----LKRKLSAET--AT----ESGFSDSGVKVIVRMK   99
PAKRP1L      HRSSPAKLKSPLP--PRPPSSNP----LKRKLIAEA--TADNGVAIGVSDSGVKVIVRMK   104

Nt-KIN12A    PANGNESGSQAVRKVSDTSVYVADRKFNFDMVFDSNSTQEDIFQSVGAPLVKDALAGYNT   162
PAKRP1       PLNKGEEGDMIVEKMSKDSLTVSGQTFTFDSIANPESTQEQMFQLVGAPLVENCLSGFNS   159
PAKRP1L      PPSKGEEEEMIVKKISNDALTINEQTFTFDSIADPESTQDEIFQLVGAPLVENCLAGFNS   164

Nt-KIN12A    SLLAYGQTGSGKTYTMWGPPSSMVEVPSPIGLQGIVPRIFQTLFSSIQREQENSEGKQIN   222
PAKRP1       SVFAYGQTGSGKTYTMWGPANGLLEEHLCGDQRGLTPRVFERLFARIKEEQVKHAERQLN   219
PAKRP1L      SVFAYGQTGSGKTYTMWGPANGLLEEHLSGDQRGLTPRVFELLFARLSEEQAKHAERQLK   224

Nt-KIN12A    YQCRCSFLEIYEEHIGDLLDPTQRNLKIMDDPRVGFYVENLTEEYVSTYEDVTQILIKGL   282
PAKRP1       YQCRCSLLEIYNEQITDLLDPSQKNLMIREDVKSGVYVENLTEEYVKNLTDVSQLLIKGL   279
PAKRP1L      YQCRCSFLEIYNEQITDLLDPSLKNLMIREDVKSGVYVENLTEEYVKNLKDLSKLLVKGL   284

Nt-KIN12A    SSRKVGSTNINSKSSRSHIVFTCIIESWCKESSSKCFGSSKMSRMNLVDLAGFERFIPDD   342
PAKRP1       GNRRTGATSVNTESSRSHCVFTCVVESRCKNVAD-GLSSFKTSRINLVDLAGSERQKSTG   338
PAKRP1L      ANRRTGATSVNAESSRSHCVFTCVVESHCKSVAD-GLSSFKTSRINLVDLAGSERQKLTG   343

Nt-KIN12A    ASKLFVKEGKHVKKSTSQLGRLVNVLSERSQSGKFDDVSYSSSALTHLMRESLGGNAKLA   402
PAKRP1       AAGERLKEAGNINRSLSQLGNLINILAEISQTGKPRHIPYRDSRLTFLLQESLGGNAKLA   398
PAKRP1L      AAGDRLKEAGNINRSLSQLGNLINILAEISQTGKQRHIPYRDSRLTFLLQESLGGNAKLA   403

Nt-KIN12A    VICAVTPENKHNSETISTLRFGKRVKLMPNEPLVNEISEDDVNGLSDQIRQLKEELIRAR   462
PAKRP1       MVCAVSPSQSCRSETFSTLRFAQRAKAIQNKAVVNEVMQDDVNFLRGVIHQLRDELQRMK   458
PAKRP1L      MVCAVSPSQSCRSETFSTLRFAQRAKAIQNKAIVNEVMQDDVNFLREVIRQLRDELQRVK   463

Nt-KIN12A    SSTSISV-GS-NGYFRGPNVRESLNQLRV-SLNRSLILPSIDNDREEEIHINEDDIKELQ   519
PAKRP1       N-DGNNPTNPNVAYSTAWNARRSLNLLRSFGLGHPRSLPHEDNDGDIEMEIDEAAVERLC   517
PAKRP1L      DDKGNNPTNPNAAYTTSWNARRSLSLLRSFGLGHPKSLPNGDDDGDTEMEIDEEAVERLC   523

Nt-KIN12     LQIDNLRGSHGDNSKETLEKKDSLKYS------------------SGESE---HYLSCSE   558
PAKRP1       VQVGLQSSLASEGINHDMNRVKSIHSSDGQS---------IEKRLPEDSDVAMEDACCHT   568
PAKRP1L      AQMGLSPPAE--DNNQEMSRVEKINSSLQTVVLKDESYNNSHLKSSEATDVNMEDACCQT   581

Nt-KIN12A    ESGNEEIN------------------------SEE----------------------TLE   572
PAKRP1       ENHEPETVDNMRTETETGIRENQIKTHSQTLDHESSFQPLSVKDALCS---------SLN   619
PAKRP1L      ENNGSETDNALT--------------VAETMDDGSSVQPDSITNSLHSCISDTNQGNSPS   627

Nt-KIN12A    ET-------------QNN-----AD---QEMEIMQPEYCNSISISPRRGSAVLQGPVLSE   611
PAKRP1       KSEDVSSCPDLVPQDVTSANVLIADGVDDP-EHLVNSASPSLCIDPVGATPVLKSPTLSV   678
PAKRP1L      KAENIPSCQDLVIEADVSAIVSVADTSNNTEQVSVNPVSPCLSVAPVSVSPVLIPPTESA   687

Nt-KIN12A    SPKFRSTQRKSLIVSSEDDIQ--C-----------SSKSPELSPLPQGDLVQSSLRSSRI   658
PAKRP1       SPTIRNSRKS-LKTSELSTASQKDSEGEN-LVTEAADPSPATSKKMNNCSSALSTQKSKV   736
PAKRP1L      SPKIRNSRKS-LRTTSMSTASQKDIERANQLTPEVVEPSPAMSTEVLNLYSALSTKKSEA   746

Nt-KIN12A    FPGPTESLAASLHRGLEIIDYHQRNSASNRSLVSFSFEHLAVKPSPLSNDKANSSVQTSS   718
PAKRP1       FPVRTERLASSLHKGIKLLESYCQSTAQRRSTYRFSFKAPDSEPSTSIS-KADAGVQTIP   795
PAKRP1L      FPVPTRQLAASLHRGMKLLDSYRQSTALRRSTFRLSYKALECKPSTVLS-KADVGVQTYP   805

Nt-KIN12A    AEGLTSRFLATSFVCPKCNRKATSSSDVQGSLRTSWMVPVEG-----ASSDQVPEDPEKV   773
PAKRP1       GADAISEENTKEFLCCKCKCREQFDAQQMGDMPNLQLVPVDNSEVAEKSKNQVPKAVEKV   855
PAKRP1L      QADEIAEDNSKEVLCSRCKCRAECDAQEISDTSNLQLVPIDNSEGSEKSNFQVPKAVEKV   865

Nt-KIN12A    LFQALEREKQLESVCKDQADKIEQLNQRLAQCKCTIEPSSLIEC-------------GKV   820
PAKRP1       LAGSIRREMALEEFCTKQASEITQLNRLVQQYKHERECNAIIGQTREDKIIRLESLMDGV   915
PAKRP1L      LAGSIRREMAMEEFCTKQASEISQLNRLVQQYKHERECNAIIGQTREDKIVRLESLMDGV   925

Nt-KIN12A    VEFKDYEKQASIIYQNGSQSLNNPKLLKWDDEESPDPEAVKEKYEIKEIQGDVDHCGGKR   880
PAKRP1       LSKEDFLDEE------FASLLHEHKLLK--DMYQNHPEVLKTKIELERTQEEVEN-FKNF   966
PAKRP1L      LSKDDFLDEE------FASLMHEHKLLK--DMYENHPEVLQTRIELKRVQEELES-FKNF   976

Nt-KIN12A    LFDMAERETLLKEIEGLRSQLQSHNGASTNKSIERTRSSLLAQSMQLRKSGA-----Y--   933
PAKRP1       YGDMGEREVLLEEIQDLKLQLQCYIDPSLKSA--LKTCTLLKLSYQ-----APPVNAIPE   1019
PAKRP1L      YGDMGEREVLLEEIHDLKAQLQCYTDSSLTSA--RRRGSLLKLTYACDPNQAPQLNTIPE   1034

Nt-KIN12A    --PKSSGEELEKERERWTEMESEWICLTDELRIDLEAHRRRAEKVAMELMLEKKCTEELD   991
PAKRP1       SQDESLEKTLEQERLCWTEAETKWISLSEELRTELEASKALINKQKHELEIEKRCGEELK   1079
PAKRP1L      SVDEGPEKTLEQERLRWTEAESNWISLAEELRTELDTNRLLMEKQKRELDTEKRCAEELT   1094

Nt-KIN12A    DALKRSVLGQARMIEHYAELQDKYNDLAEKHKLILQGIQDVKMAAAKAGRKGHGARFAKS   1051
PAKRP1       EAMQMAMEGHARMLEQYADLEEKHMQLLARHRRIQDGIDDVKKAAARAGVRGAESRFINA   1139
PAKRP1L      EAMQMAMQGHARMIEQYADLEEKHIQLLARHRRIREGIDDVKKAAARAGVKGAESRFINA   1154

Nt-KIN12A    LAAELSALRVERERERERMLKKENRSLRAQLKDTAEAVHAAGELLVRLREAEETASVAEEN   1111
PAKRP1       LAAEISALKVEKEKERQYLRDENKSLQTQLRDTAEAIQAAGELLVRLKEAEEGLTVAQKR   1199
PAKRP1L      LAAEISALKVQREKEVRYFRDENKSLQSQLRDTAEAVQAAGELLVRFKEAEEGLTFAQKR   1214

Nt-KIN12A    FTKSKEENEKLKKQIEKLKRKHKMEMITMKQYLAESR-LPEAALRPPLYREDSDVANSET   1170
PAKRP1       AMDAEYEAAEAYRQIDKLKKKHENEINTLNQLVP------QSHIH----NECSTKCDQAV   1249
PAKRP1L      AMDAEYEASEAYKKVDKLKRKYETEISTVNQQHNAEPQNPIESLQ----ASCNDDAMAKY   1270

Nt-KIN12A    VQHHEYDDDQAWRAEFGAIYQERI-------------------          1194
PAKRP1       EPSVNASSEQQWRDEFEPLYKKETEFSNLAEPSWFSGYDRCNI          1292
PAKRP1L      DEPSASDGDNQWREEFQPFYKKDEELSKLAEPSWFSGYDRCNI          1313
```

**Fig. S5.** Alignment of the amino acid sequence of Nt-KIN12A in *N. tabacum* with the presumptive homologs in Arabidopsis.

The proteins Nt-KIN12A (GenBank ID: NP_001312269.1) from *N. tabacum* and PAKRP1 (AT4G14150) and PAKRP1L (AT3G23670) from Arabidopsis. The similarity (identity) of the full-length amino acid sequence of Nt-KIN12A with each Arabidopsis protein was 48% and 45%, respectively. The identified phosphorylated residues are shown in red with the arrowheads. The putative kinesin motor domain and coiled-coil region are colored in yellow and cyan, respectively.

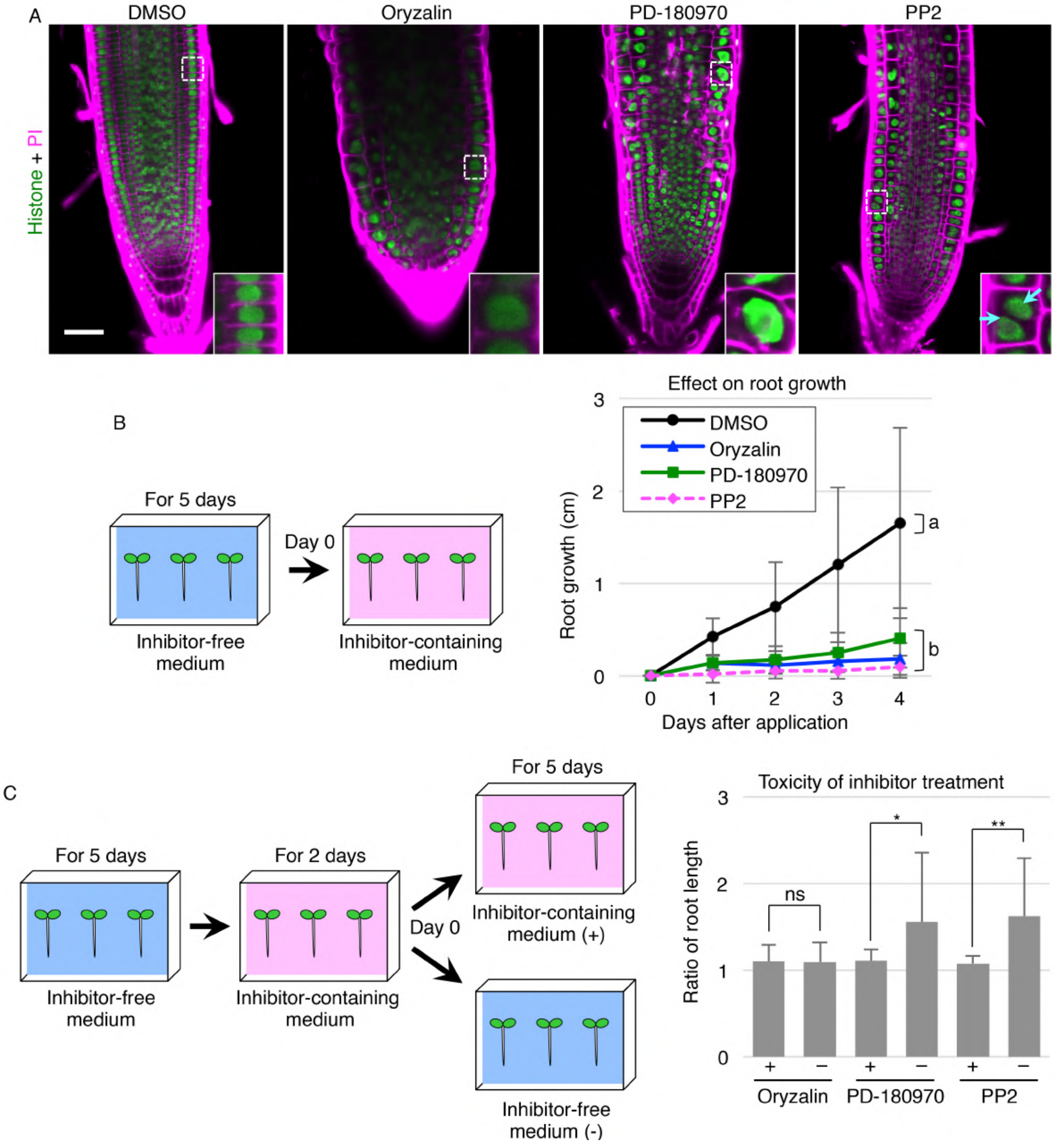


**Fig. S6.** Effects of PD-180970 and PP2 on Arabidopsis root.

(A) Confocal images of PI-stained root tips in 6-day-old Arabidopsis seedlings expressing histone marker. The roots were observed at 1 day after the transferring onto the media containing indicated compounds. Insets show the enlarged images of the dotted square areas, and cyan arrowheads point two nuclei in a cell. (B) Root growth after the application of indicated compounds. Graph shows the increase from the root length at day 0, and error bars represent SD ($n \geq 21$). The letters on the graph indicate significant differences of the values at 4 days after application, which were determined by the Tukey–Kramer test ($P < 0.01$). The left illustration shows the schematic procedure of the experiment. (C) Toxicity of inhibitor treatment on Arabidopsis root growth. As shown in the left illustration, 5-day-old seedlings germinated on inhibitor-free media were then grown on the media containing indicated compound for 2 days, and transferred to a new media containing the same compound (+) or no reagent (-). Graph shows the ratio of root length at 5 days to 0 days after the final transference. Error bars represent SD ($n \geq 19$). Significant difference was determined by Brunner–Munzel test ($P < 0.01$ (**), $P < 0.05$ (*), or ns (not significant)). Scale bar: 50 μm.

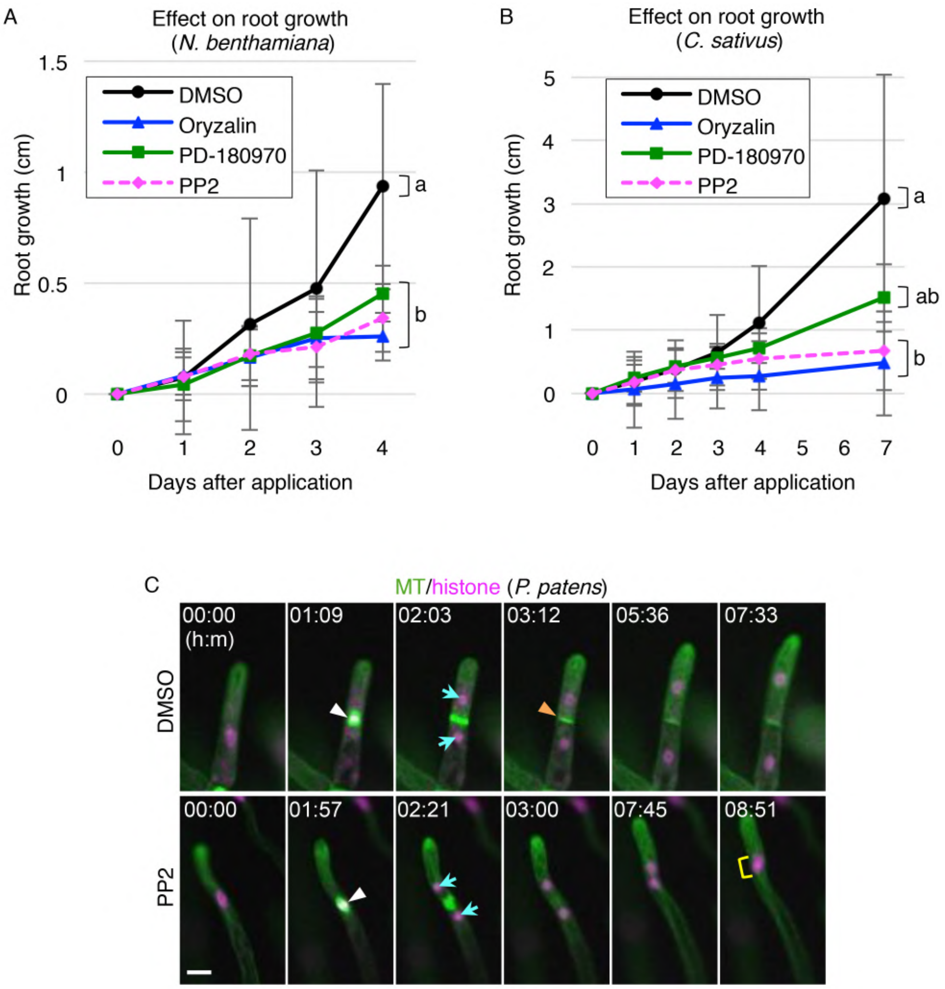


**Fig. S7.** Effects of PD-180970 and PP2 on various plant species

(A and B) Root growth after the application of indicated compounds to 5-day-old seedlings of *N. benthamiana* (A) and *C. sativus* (B). Graph shows the increase from the root length at day 0, and error bars represent SD (n ≥ 9 (A), and 8 (B)). The letters on the graph indicate significant differences of the values at 4 days (A) or 7 days (B) after application, which were determined by the Tukey–Kramer test ($P < 0.01$). (C) Time-lapse observation of *P. patens* chloronema cells expressing MT/histone marker in the presence of indicated compounds. Numbers indicate the time (h:min) from the first frame. White and orange arrowheads indicate the condensed nuclei and newly formed cell plate, respectively. Cyan arrows and yellow rectangle show separated nuclei and two accompanied nuclei, respectively. Scale bar: 20 μm.

**Supplementary Table**

**Table S1.** Quantification of the inhibitory effects in tobacco BY-2 cells.

| | | | Appearance of abnormal cell division (%) | P-value (vs. DMSO) | n |
|---|---|---|---|---|---|
| Related compounds | | DMSO | 3 | | 60 |
| | | oryzalin | 100 | 0 | 36 |
| | PD-180970 -related | PD-180970 | 100 | 0 | 44 |
| | | PD-166326 | 100 | 0 | 27 |
| | | PD-173955-Analog1 | 13 | 0.1 | 62 |
| | | Ponatinib | 70 | 0 | 27 |
| | | Bosutinib | 17 | 0.03 | 29 |
| | | Bafetinib | 12 | 0.15 | 25 |
| | PP2-related | PP2 | 100 | 0 | 55 |
| | | PP1 | 100 | 0 | 32 |
| | | PP2-Analog1 | 0 | 1 | 13 |
| | | PP2-Analog2 | 0 | 0.54 | 33 |
| | | PP3 | 0 | 1 | 20 |
| | | Src inhibitor1 | 5 | 0.67 | 55 |

Quantification of the inhibitory effects in tobacco BY-2 cells.
The appearance of abnormal cell division shows the ratio of BY-GTRC cells that initiated cell division but did not complete it. P-values were calculated using two-tailed Fisher's exact test.

**Supplementary Movies**

**Movie S1 (separate file).** Effect of PD-180970 and PP2 on zygotic cell division in Arabidopsis.

Time-lapse observation of zygotes expressing histone/PM markers in the presence of the indicated compounds. MIP images are shown. Images were obtained at 10-min intervals, and the numbers indicate the time (h:min) from the first frame. Scale bar: 10 μm.

**Movie S2 (separate file).** Effects of PD-180970 and PP2 on MT organization in Arabidopsis zygotes.

Time-lapse observation of Arabidopsis zygotes expressing MT markers in the presence of the indicated compounds. MIP images are shown. Images were obtained at 20-min intervals, and the numbers indicate the time (h:min) from the first frame. Scale bar: 10 μm.

**Movie S3 (separate file).** Effects of PD-180970 and PP2 on tobacco BY-2 cells.

Time-lapse observation of BY-GTRC in the presence of the indicated compounds. MIP images are shown. Images were obtained at 10-min intervals, and the numbers indicate the time (h:min) from the first frame. Scale bar: 100 μm.

**Movie S4 (separate file).** Effect of PP2 on PAKRP1 localization and phragmoplast formation in tobacco BY-2 cells.

Time-lapse observation of BY-2 cells expressing the PAKRP1/MT marker in the presence or absence of PP2. Images were obtained at 30-sec intervals, and numbers indicate the time (min: s) from the onset of anaphase. Scale bar: 10 μm.

**Movie S5 (separate file).** Effect of PP2 on PAKRP1L localization and phragmoplast formation in tobacco BY-2 cells.

Time-lapse observation of BY-2 cells expressing the PAKRP1L/MT marker in the absence of compounds and PP2. Images were obtained at 30-sec intervals, and the numbers indicate the time (h:min) from anaphase onset. Scale bar: 10 μm.

**Movie S6 (separate file).** Effect of PP2 on class II Kinesin-12 localization and phragmoplast formation in *P. patens* chloronema.

Time-lapse observation of *P. patens* chloronema expressing Pp-Kinesin-12IIc/MT marker in the presence of the indicated compounds. Images were obtained at 30-sec intervals, and numbers indicate the time (min: s) from the onset of anaphase. Scale bar: 10 μm.

**Movie S7 (separate file).** The effect of PP2 on cytokinesis in *P. patens* chloronema.

Time-lapse observation of *P. patens* chloronema expressing MT/histone markers in the presence of the indicated compounds. Images were obtained at 3-min intervals, and the numbers indicate the time (h:min) from the first frame. Scale bar: 100 μm.

**Supplementary Datasets**

**Dataset S1 (separate file).** Candidates of PD-180970 phosphorylation target.

Top candidates obtained from phosphoproteomics of BY-GTRC cells, which were synchronized at the M phase in the presence of the effective reagent (PD-180970) and ineffective control (PD-173955-Analog1). The peptide number shows the count of identified phosphopeptides in the 1st and 2nd experiments. The ratio shows the value of the peptide number of PD-180970 divided by that of the PD-173955-Analog1 for each experiment. This table shows the proteins whose peptide number was 10 or more, and whose ratio was less than 0.5 in both experiments, as the top candidates of the phosphorylation target of PD-180970. Reference ID shows the code in our database generated from RNA-seq data, and its corresponding protein in Genbank is shown with their identity.

**Dataset S2 (separate file).** Candidates of PP2 phosphorylation target.

Top candidates obtained from the phosphoproteomics of BY-GTRC cells, which were synchronized at the M phase in the presence of the effective reagent (PP2) and the ineffective control (PP3). The peptide number shows the count of identified phosphopeptides in the 1st, 2nd, and 3rd experiments. The ratio shows the value of the PP2 peptide number divided by that of PP3 for each experiment. This table shows the proteins whose peptide number was 10 or more in all experiments, and whose ratio was less than 0.5 at least twice in 3 experiments, as the top candidates of the PP2 phosphorylation target. Reference ID shows the code in our database generated from RNA-seq data, and its corresponding protein in Genbank is shown with their identity.
